# Androgen receptor signaling blockade enhances NK cell-mediated killing of prostate cancer cells and sensitivity to NK cell checkpoint blockade

**DOI:** 10.1101/2023.11.15.567201

**Authors:** Maximilian Pinho-Schwermann, Connor Purcell, Lindsey Carlsen, Kelsey E. Huntington, Praveen Srinivasan, Andrew George, Vida Tajiknia, William McDonald, Lanlan Zhou, Leiqing Zhang, Andre De Souza, Howard P. Safran, Benedito A. Carneiro, Wafik S. El-Deiry

## Abstract

**Background:** The blockade of the androgen receptor (AR) pathway is an effective treatment for prostate cancer (PCa), but many patients progress to metastatic castration-resistant prostate cancer (mCRPC). Treatments for mCRPC include AR inhibitors (ARi), chemotherapy, PARP inhibitors, and radioligands. Checkpoint inhibitor activity is limited to a small subset of MSI-H mCRPC. AR signaling modulates CD8+ T cell function, but its impact on natural killer (NK) cell cytotoxicity is unknown. We investigated the effect of ARi on NK cell activation, cytokine secretion, NKG2A expression, and NK cell-mediated killing of PCa cells in vitro.

**Methods:** PCa cell lines (LNCaP, 22Rv1, DU145, PC3) were co-cultured with NK-92 and treated with ARi (enzalutamide [enza], darolutamide [daro]) alone or in combination with anti-NKG2A antibody monalizumab. Immune cell-mediated tumor cell killing and cytokine secretion were quantified. NK cell expression of NKG2A and PCa cell expression of HLA-E were investigated by flow cytometry. The AR-negative cell lines PC3 and DU145 were stably transduced with an AR expression vector to evaluate the AR modulation of HLA-E. To assess the in vivo combination of NKG2A blockade and ARi therapy in vivo, Cas9 was used to genetically ablate the murine HLA-E ortholog, H2-T23, from RM-1 murine PCa cells. H2-T23 knockout and control cells were grown subcutaneously in castrated C57BL/6 mice and treated with daro or control. The activation status of peripheral blood NK cell isolated from patients with PCa before and after initiation of androgen deprivation therapy (ADT) was evaluated by flow cytometry.

**Results:** ARi activated NK cells and significantly increased immune-mediated NK-92 cell killing of PCa cells. IFN-γ and TRAIL mediated ARi-induced NK cell activation. ARi increased expression of the inhibitory receptor NKG2A on NK cells, and immune killing of PCa cells was enhanced with the combination of ARi and monalizumab. ARi also increased the expression of HLA-E, the ligand of NKG2A, on PCa cell lines. By transducing AR into AR-negative PC3 and DU145, we demonstrated that androgen signaling regulates HLA-E expression. In a mouse model of PCa, HLA-E knockout synergized with darolutamide to increase NK cell activation. NK cells derived from patients with metastatic PCa exhibited increased expression of Granzyme B and Perforin following ARi treatment.

**Conclusions:** ARi activates NK cells via IFN-γ and TRAIL and promotes the killing of PCa cells. ARi also upregulates expression of HLA-E on PCa which may suppress the innate immune response against PCa. ARi-mediated NK cell killing of PCa cells was enhanced by NKG2A blockade. These results support novel immunotherapeutic strategies for PCa targeting NK activation through the combination of ARi and monalizumab.

**Graphical Abstract:** Androgen receptor inhibitors (ARi) enhance NK cell-mediated killing of prostate cancer cells and sensitivity to NK cell checkpoint NKG2A blockade. ARi upregulate the NK cell inhibitor ligand (HLA-E) mediating suppression NK cell killing of PCa. This regulation is dependent on a functional AR signal on tumor cell lines. Adding an anti-NKG2a-HLA-E mAb with ARi further enhances the NK cell-mediated killing of PCa.

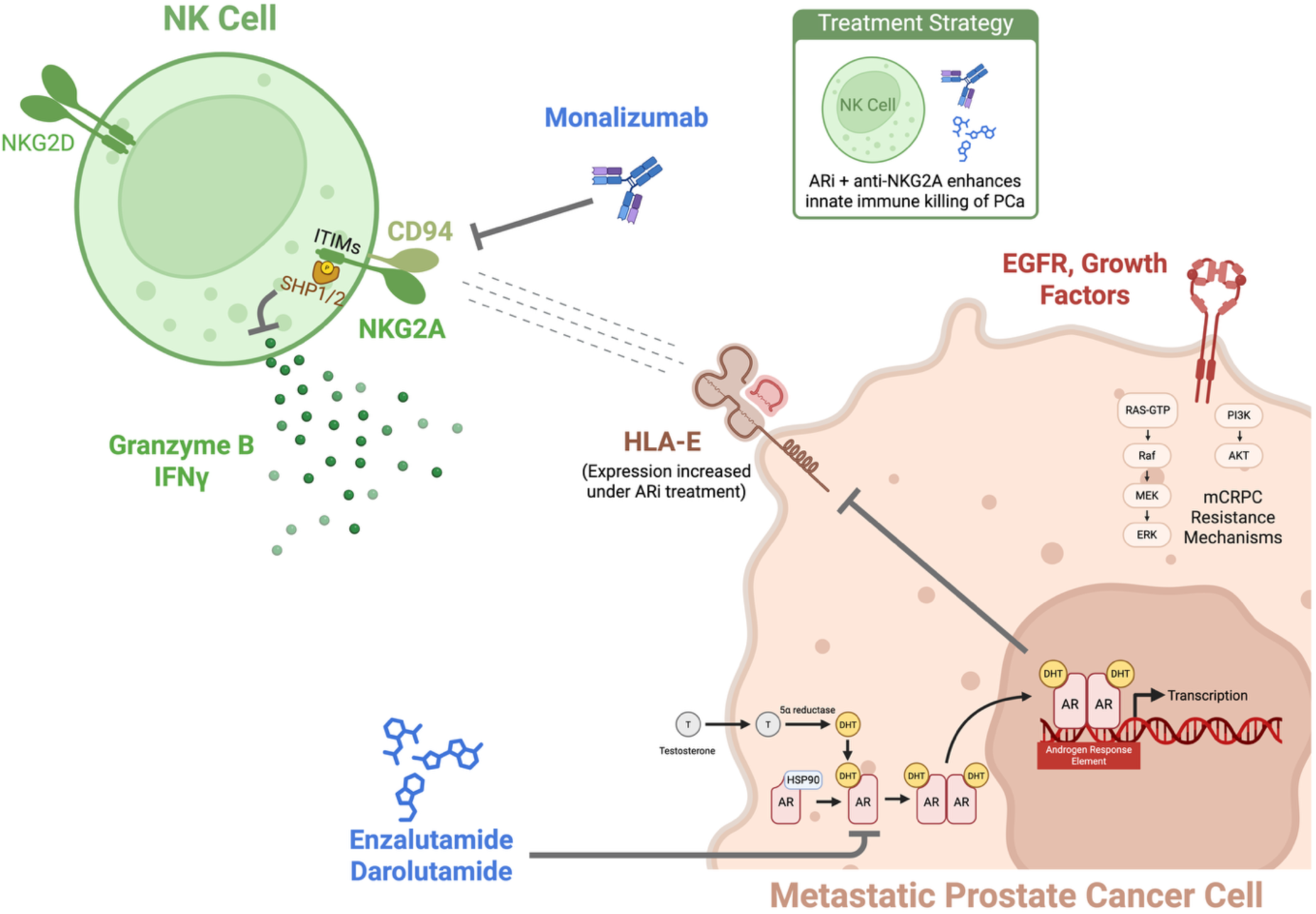

## Introduction

Prostate cancer (PCa) is the most common cancer diagnosed in men, ranking second in cancer-related mortality in the United States [1]. Although most patients present with localized disease, progression to metastatic disease and its management remain a significant clinical challenge. Blockade of androgen receptor (AR) signaling is central to PCa treatment; however, most patients with advanced disease progress to metastatic castration-resistant prostate cancer (mCRPC), which is associated with a median OS of 4-5 years [2]. Standard treatments for mCRPC include AR-targeted agents (e.g. enzalutamide, darolutamide, apalutamide), the CYP17 inhibitor abiraterone, chemotherapy (e.g. docetaxel, cabazitaxel), Lutetium 177-PSMA-617, and PARP inhibitors. Although the first therapeutic cancer vaccine targeting prostate tumors was approved more than a decade ago (sipuleucel-T), the clinical efficacy of immunotherapy for treating PCa remains limited, with poor responses to checkpoint inhibitors [3]. PCa tumors display an immunosuppressive tumor microenvironment mediated by reduced expression of human leukocyte antigen (HLA) molecules, decreased neoantigen expression, phosphatase and tensin homolog (PTEN) protein loss, and dysfunction of type I interferon (IFN) signaling [4, 5]. These factors contribute to low response rates to checkpoint inhibitors in most cases of mCRPC, except for microsatellite instability-high (MSI-H) tumors and those with CDK12 mutations [6, 7]. Accordingly, novel immunotherapy strategies, including T cell engagers, represent an active area of research to overcome the immunologically “cold” PCa phenotype.

Androgen signaling modulates the immune response by inhibiting CD8 and CD4 T cells [8]. AR blockade enhances CD8+ T cell function and sensitizes tumors to PD-1 checkpoint inhibitors in animal models of prostate cancer [8, 9]. Androgen signaling in T cells suppresses IFN-γ secretion in vitro and contributes to T cell exhaustion. AR blockade reversed these effects and contributed to PSA and tumor responses among patients with mCRPC treated with pembrolizumab (anti-PD-1) and enzalutamide in a clinical trial [10]. In addition, gene expression analysis of CD4+ T cells isolated from castrated mice identified protein tyrosine phosphatase non-receptor type 1 (Ptpn1) as a mediator of androgen-induced suppression of CD4+ T-cell differentiation [11–13]. Upregulation of interferon regulatory factor-1, 3, and 7 (IRF-1,3 and 7) and STAT4 in CD4+ T cells from mice were also observed, suggesting androgens may alter IL-12-induced STAT4 phosphorylation and impair CD4+ T cells. While these results highlight the role of AR signaling in CD8 and CD4 T cells, the impact on natural killer (NK) cells has yet to be elucidated.

In this study, we investigated the effect of AR inhibitors enzalutamide (enza) and darolutamide (daro) on NK activation *in vitro* using co-culture experiments with immune cells, PCa cell lines, patient-derived immune cells, and a syngeneic prostate cancer animal model. The results demonstrated that AR inhibitors activate NK cells and enhance the immune-mediated killing of PCa cells through an IFN-γ- and TRAIL-mediated mechanism. This effect was potentiated by the combination with monalizumab, which blocks the immunosuppressive NK cell receptor NKG2A. We also identified the androgen-dependent modulation of HLA-E in PCa cells as a potential mechanism for suppressing the innate immune response and protecting PCa cells from NK cell killing. These results support the development of novel immunotherapy approaches in prostate cancer that promote NK cell activation through AR inhibition and NKG2A blockade, among other potential NK cell-based therapeutics.

## Materials and Methods

### Cell lines and culture conditions

Human prostate cancer cell lines (obtained from ATCC) included PC3, DU145, LNCaP, and 22Rv1. Murine RM-1 cells were a gift from Tejal Desai, previously obtained from ATCC. Cells were grown in RPMI media supplemented with 10% FBS, 1% sodium pyruvate, 1% Glutamax, and 1% penicillin/streptomycin at 37°C, 5% CO_2_. Cells were trypsinized (0.25% trypsin) upon flask confluency and tested for mycoplasma every 6 months. Even though 22Rv1 constitutively expresses the androgen receptor, it is resistant to androgen blockade due to a splice-site mutation (AR-V7) [14].

NK cell line NK-92 was obtained from ATCC. The cell line was grown in alpha-MEM medium supplemented with 10% FBS, 1% sodium pyruvate, 1% Glutamax, 1% penicillin/ streptomycin, 0.1 mM of 2-mercaptoethanol, 12.5% of heat-inactivated horse serum, 1% Non-Essential Amino Acids (NEAA), 1% folic acid and 20 mM myoinositol at 37°C, 5% CO_2_. NK-92 cells were kept at 4-5 x 10^5^ cells/ml seeding concentration and supplemented with 100 IU/ml of recombinant human IL-2 every two days. Cells were passaged using Ficoll PM400, sodium diatrizoate, and disodium calcium EDTA solution (Ficoll-Paque PLUS Cytiva 17-1440-02) every 10-14 days and up to 72 h before plating/experiments.

### Western blot analysis

Androgen receptor expression in prostate cancer cell lines and immune cells (NK-92) was examined using a western blot. A total of 1×10^5^ cells were plated and incubated in a 6 or 12-well plate (CELLTREAT 229111 and 229105) for 24 hours to allow the cells to adhere (tumor cells). Treatment conditions were added after cell adhesion or in conjunction with NK cells when these were analyzed. Cells were harvested and lysed using RIPA buffer containing protease inhibitor (Cell Signaling Technology 9806S) and phosphatase inhibitor (PHOSS-RO Roche 4906845001). Denaturing sample buffer was added, samples were boiled at 95°C for 10 minutes, and an equal amount of protein lysate (10-20 μg) was electrophoresed through 4–12% SDS-PAGE gels (Invitrogen) and then transferred to PVDF membranes. The membrane was blocked with 5% milk in 1x TTBS and incubated overnight with the appropriate primary antibody (Cell Signaling Ran Antibody #4462; Sigma Aldrich Monoclonal Anti-β-Actin antibody #A5441; Cell Signaling Androgen Receptor (E3S4N) Rabbit mAb (Carboxy-terminal Antigen) #70317; Cell Signaling Vinculin antibody #4650; Invitrogen QA-1b Monoclonal Antibody #MA5-24801). The primary antibody membranes were incubated in the appropriate HRP-conjugated secondary antibody (either mouse, Thermo Scientific #31430, or rabbit, Thermo Scientific #31460) for two hours. The levels of antibody binding were detected using ECL western blotting detection reagent and the Syngene imaging system.

### Establishing IC50 doses

PCa cells were plated at a density of 5 x 10^3^ cells per well of a 96-well plate. Cells were treated with doses ranging from 0-250 µM of anti-androgens Enzalutamide and Darolutamide. 5 x 10^3^ cells were plated in a 96-well plate Cell for the viability assays requiring NK cells, and viability (72 h) was measured using a CellTiterGlo assay. Bioluminescence imaging was measured using the Xenogen IVIS imager. IC50 doses were determined based on the dose-response curve collected from the data. Concentrations used in the experiments were correlated with those findings, matched with literature reports, and clinical correlation when possible.

### Co-culture assays

PCa cell lines were stained with the blue CMAC live-cell dye (Thermo Fisher Scientific #C2110). Blue-fluorescent cancer cells were plated at a density of 10,000 cells per well of a 48-well plate. NK-92 cells were dyed with green CMFDA (Cayman Chemical Company #19583). Green-fluorescent NK-92 cells were added to the blue-fluorescent cancer cells at 1:1. Enzalutamide and 1 µM red fluorescent ethidium homodimer (EthD-1) were added at the same time as the NK cells. After 24 hours of co-culture, images of live cancer cells, live NK cells, and dead cells were taken using a fluorescent microscope. The co-culture experiments were run with internal control conditions and as a single treatment. Normalization was performed after analysis to take baseline death conditions into each experiment run. To optimize the high output of these experiments, a subset of co-cultures was performed using the ImageXpress® Micro Confocal High-Content Imaging. This analysis maintained the experiment conditions, drug concentrations, and the dying process. When GFP-labeled tumor cells were used for culture assays, they were added in a 1:1 ratio. Cell adhered over 24 h, and when patient-derived NK cells were added, a control well was trypsinized, and cells resuspended and counted to adjust a 1:1 ratio before treatment addition. The experiments with blocking antibodies were performed using the ImageXpress system. TRAIL blocking ab was purchased from Santa Cruz TRAIL Antibody (RIK-2): sc-56246, and IFN-γ–blocking Monoclonal Antibody (NIB42) was purchased from Thermo Fisher. The blocking mAb was added when NK cells were combined with tumor cells.

### Mice/Tumor Models

To generate knockout cells, Alt-R™ S.p. Cas9-GFP V3 recombinant Cas9 protein (IDT 10008100) was complexed with nontargeting or H2-T23 sgRNA (Genscript) according to the manufacturer protocol and delivered to RM-1 prostate cancer cells with Lipofectamine™ CRISPRMAX™ transfection reagent (ThermoFisher CMAX00008). The media was replaced 48hr later and cells were cultured. Protein expression was assessed by western blot.

C57BL/6J (Jackson #000664) mice (n=8 per group) were purchased from Jackson Laboratories. All mice were maintained under pathogen-free conditions in the laboratories of pathology and biology at Brown University, Providence, RI. All animal studies were approved and performed according to the Center for Animal Care Procedures (CARE) at Brown University and Institutional Animal Care and Use Committee (IACUC) protocols. A total of 1 × 10^6^ RM-1 cells were suspended in 100 μl of ice-cold PBS and 100 μl of Matrigel® Basement Membrane Matrix (Corning® #354234). Anesthesia using 100 mg/kg ketamine was performed before a total of 200μl cell suspension was inoculated subcutaneously into the rear left flank of the mice aged 6-8 weeks. Male mice were assigned to a treatment group. Upon randomization, all mice received androgen-deprivation therapy using 0.5 µg degarelix injected subcutaneously. On the same day, darolutamide (50 mg/kg, oral gavage, twice daily) or vehicle was delivered in a solution of 40% PEG400, 5% Tween 80, 5% DMSO and 5% PBS. The experiment was continued until the tumor burden had reached the ethically allowed limit at 2000 mm^3^ in volume or when the mouse had lost more than 20% of its initial body weight. At the end of the experiment, mice were sacrificed per Brown University CARE and IACUC protocols. Tumors and organs were harvested for immunohistochemistry.

### Statistical analysis

Live cancer cells, NK cells, and dead cells were quantified using FIJI (ImageJ) software for images derived from fluorescent co-culture images. ImageXpress images were quantified using its software analysis of cytoplasmic/nuclear staining. The percent of dead tumor cells in each well was quantified, and this percentage was normalized by subtracting baseline death from cancer cell-only wells, NK cell-only wells, or both. A two-way ANOVA was used to calculate the interaction effect between drug treatment and NK cells.

In groups with a significant interaction effect, data was further normalized in the cancer cell + NK cells + drug treatment well by subtracting out death in the cancer cell + drug well. A one-way ANOVA and multiple comparison analysis with subsequent t-tests were used to calculate the statistical significance of the difference between this group and the cancer cell + NK cell well.

### Generation of stably expressing GFP, AR, and AR Activity Reporter cell lines

#### GFP-Positive Cell Line Production

The prostate cancer cell lines (PC3, DU145, LNCaP, and 22Rv1) were seeded at 50% confluence in a 12-well tissue culture plate and adhered overnight. Then, they were transduced with lentivirus containing pLenti_CMV_GFP_Hygro [pLenti CMV GFP Hygro (656-4) was a gift from Eric Campeau and Paul Kaufman (Addgene viral prep #17446-LV)] at a multiplicity of infection of 10 for 48 hours before washing with PBS and replacing with fresh medium. The cells were then sorted for GFP-positivity using a BD FACSAria™ III Cell Sorter (RRID: SCR_016695).

#### Plasmid handling

Agar stabs with plasmid-containing bacteria were obtained from Addgene. Bacteria were streaked onto LB agar dishes with 100 µg/mL ampicillin and grown at 30°C for 24 hours. Single colonies were then Midi-prepped (QIAGEN Plasmid Plus Midi Kit, 12943) according to the manufacturer’s protocol. The plasmid sequence was verified by whole-plasmid sequencing (Plasmidsaurus) using Oxford Nanopore technology.

#### Lentivirus production

Five million HEK293T cells (ATCC CRL-3216) were seeded in a 10 cm tissue culture dish with 6 mL of antibiotic-free DMEM with 10% FBS (ATCC 30-2020) and adhered overnight. They were then transfected using 50 µL of Lipofectamine 2000 (Thermo Fisher 11668019), 10 µg of transfer plasmid [either pLENTI6.3/AR-GC-E2325 (Addgene 85128, pLENTI6.3/AR-GC-E2325 was a gift from Karl-Henning Kalland) or ARR3tk-eGFP/SV40-mCherry (Addgene 132360, ARR3tk-eGFP/SV40-mCherry was a gift from Charles Sawyers)], 5 µg of pMDLg/pRRE (Addgene 12251, pMDLg/pRRE was a gift from Didier Trono), 5 µg pRSV-Rev (Addgene 12253, pRSV-Rev was a gift from Didier Trono), and 2.5 µg of pMD2.G (Addgene 12259, pMD2.G was a gift from Didier Trono). After 16 hours, the medium was exchanged. Forty-eight hours after transfection, the medium was harvested, centrifuged at 500g for 5 minutes, and the supernatant was sterile-filtered through a 0.45 µm polyether sulfone syringe filter (Millipore SLHPR33RS). The supernatant was then stored at −80°C for future use.

#### Lentivirus Transduction for pLENTI6.3/AR-GC-E2325 and ARR3tk-eGFP/SV40-mCherry

PC3 and DU145 cell lines were seeded at 50% confluence in a 12-well tissue culture plate and allowed to adhere overnight. Varying volumes of viral supernatant were added to each well for 48 hours. Wells were then washed with PBS and replaced with fresh medium. For pLENTI6.3/AR-GC-E2325, cells were selected with 2.5 µg/ml (PC3) and 5 µg/m (DU145) of blasticidin (InvivoGen ant-bl-1) for 7-9 days. For ARR3tk-eGFP/SV40-mCherry, cells were sorted for mCherry-positivity using a BD FACSAria™ III Cell Sorter (RRID: SCR_016695).

### Human Isolation of NK Cells

NK cells were isolated from patients harboring metastatic castration-sensitive prostate cancer using the MojoSort™ Human NK Cell Isolation Kit (#480053). Cells were isolated from patients using a density-dependent method with Ficoll PM400, sodium diatrizoate, and disodium calcium EDTA solution (Ficoll-Paque PLUS, Cytiva 17-1440-02). Following the suggested protocol from the manufacturer of negative selection sorting, non-natural killer cells were depleted by incubating the patient’s whole PBMCs sample with the biotin antibody cocktail (BioLegend #480053) followed by incubation with magnetic Streptavidin Nanobeads. The magnetically labeled fraction (non-NK cell population) was retained using a magnetic separator (BioLegend #480019). The untouched NK cells (CD56+, CD3-) were collected, and the purity of isolation was determined by flow cytometry using anti-CD56 (BioLegend #362503). NK isolated cell population was only kept in NK cell medium (see methods above) and further plated for experiments if the purity of the sample for CD56+ was above 98%. The patient specimens were collected under the Brown University Health research protocol designed to investigate molecular and genetic features of tumors and mechanisms of resistance (Rhode Island Hospital IRB protocol number 449060-38). All patients provided written informed consent.

### Cytokine Analysis

3×10^5^ PC3, LNCaP, 22Rv1, and DU145 cells were plated per well of a 24-well plate and incubated for 72h hours before treatment with 15 nM of DHT. 2×10^5^ NK-92 and TALL-104 cells were incubated in 15 nM of DHT for 72h before enza or daro treatment. The medium was collected 24 hours after treatment and stored at −80°C until readout. Samples were shipped to Brown University to run in biological duplicate on a Luminex 200 Instrument (R&D LX200-XPON-RUO), which captures cytokines on magnetic antibody-coated beads and measures cytokine levels using a system based on the principles of flow cytometry (see https://www.luminexcorp.com/luminex-100200/#overview for more information). A custom 52-cytokine panel was split into 34-plex and 18-plex assays (R&D LXSAHM) and was run on the Luminex 200 instrument according to the manufacturer’s protocol.

### Flow Cytometry

PCa cells were plated at a density of 3 x 10^5^ cells per well of a 6-well plate and adhered overnight. In experiments where the treatment conditions included Enza and Darolutamide, cells were kept in culture in the presence of DHT (20-30 nM) for at least 72 h. After adhesion, cells were trypsinized, stained, and incubated (Cell Staining Buffer Southern Biotech #0225-01S) with corresponding antibody for 45-60 min at 4 C. Viability dye was added with each run. NK-92 cells and patient-derived cells were stained similarly. Markers used for experiments included: Thermofisher HLA-E Monoclonal Antibody (3D12HLA-E), Alexa Fluor™ 488, eBioscience™, CD274 (PD-L1, B7-H1) Monoclonal Antibody (MIH1), eFluor™ 450, eBioscience™, Cell Signaling PD-1 (D4W2J) XP® Rabbit mAb, SYTOX™ Orange Dead Cell Stain, for flow cytometry, BioLegend APC anti-human CD159a (NKG2A) Antibody #375108, BioLegend APC anti-human CD69 Antibody #310909, BioLegend Alexa Fluor® 647 anti-human CD137 (4-1BB) [4B4-1] #309823. For staining of the patient-derived PBMCs and subsequent NK cell characterization, the following antibodies were used: BioLegend Anti-Perforin Mouse Monoclonal Antibody (PerCP Cy5.5®, clone: dG9) #308113, BioLegend Anti-Granzyme B Mouse Monoclonal Antibody (FITC, clone: GB11) #515403, BioLegend Anti-CD56 Mouse Monoclonal Antibody (Alexa Fluor® 700, clone: 5.1H1) #362521, and BioLegend (PE anti-human CD45 Antibody) #304008.

## Results

### Androgen receptor inhibitors enhance NK cell killing of prostate cancer cells *in vitro*

Prostate cancer cell lines (22Rv1, LNCaP, PC3, and DU145) were plated on 48-well plates with DHT (15-30 nM), and immune cells were added 24 hr later (1:1 ratio). They were treated with enzalutamide (10 µM) for 48 hr. Doses of AR inhibitors (ARi) used in co-culture experiments did not impact NK cell viability (**Supp. Fig.1**) and were based on previous IC50 experiments [15–17].

**Figure 1.**
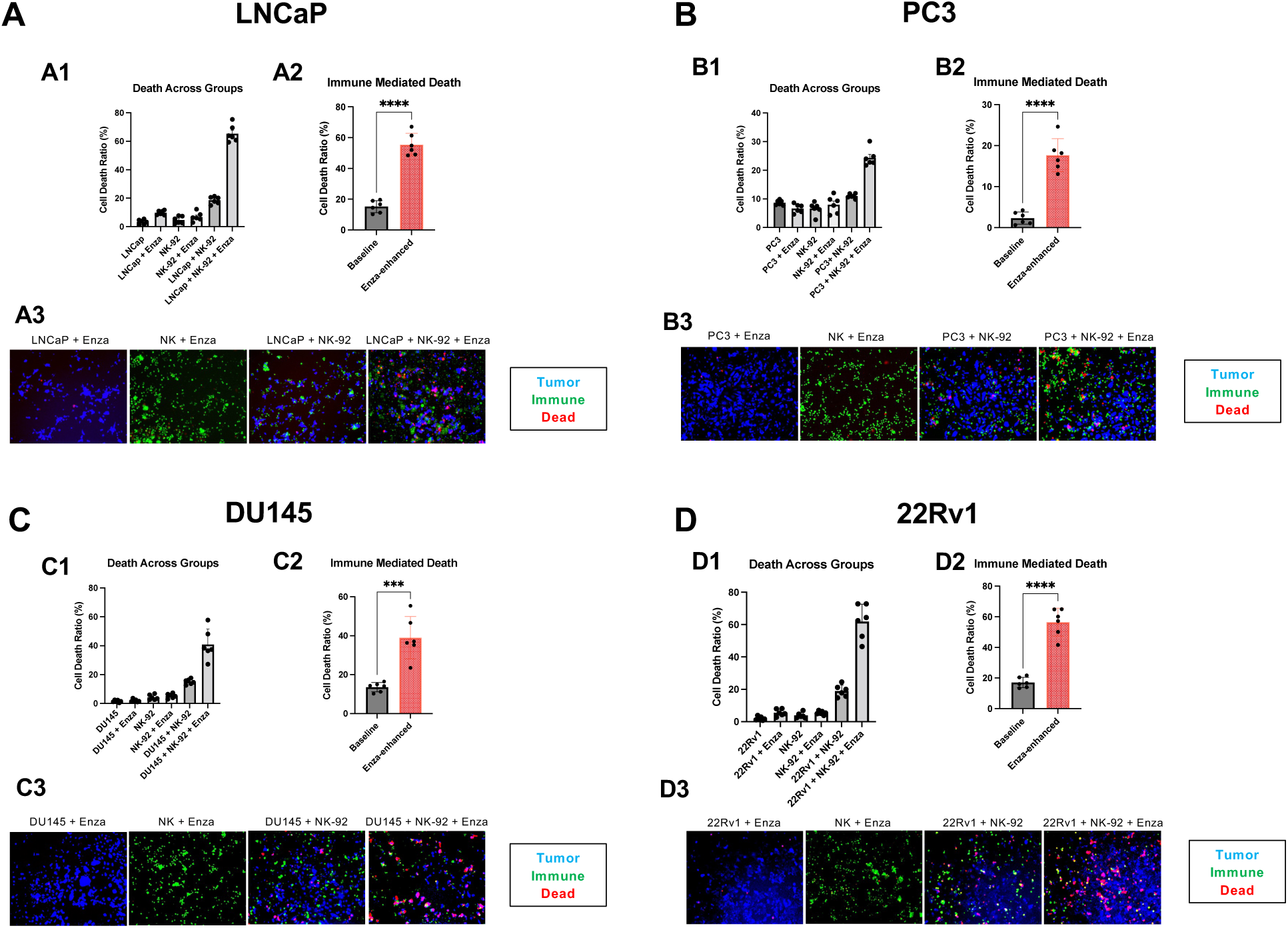
ARi enhances NK-cell killing of PCa cell lines *in vitro*. Results with the PCa cell lines (A: LNCaP, B: PC3, C: DU145, and D: 22Rv1) and the NK-92 cell line with or without enzalutamide (10 µM). Cells were cultured in a 1:1 ratio for 24 h. Cell viability was evaluated 48 h after addition of enzalutamide. Tumor cells were dyed with CMAC blue dye, and the NK-92 cell line was dyed with CMFDA green dye. The ethidium homodimer red dye (1 µM) was used to assess live dead cells. Experiments were performed with tumor cells alone and in the presence of enzalutamide to account for drug-induced cell death alone (A1, B1, C1, and D1). Quantifications were performed using Image J. Every co-culture experiment had an NK-92 control condition alone to validate the nontoxicity of enzalutamide on the immune population (data not shown). The enzalutamide dose was noncytotoxic on the NK-92 cell line (Supp. Fig. 1). The immune-mediated effect of NK-92 on the PCa cell lines in the presence of enzalutamide [Enza-enhanced = (PCa cell+NK cell+Enza) – (PCa cell + Enza)] was compared to the baseline immune-mediated effect [Baseline = (PCa cell + NK) – (PCa cell)] (A2, B2, C2, and D2). Immune enhanced killing removes the effect of enza alone from PCa cells + immune cells + enza, thus highlighting the effect of NK cell killing. The results show an increase of immune-mediated NK cell killing of PCa cell lines, irrespective of AR status or sensitivity to enzalutamide when treated with the ARi (p-value legends as follows: ns *p* > 0.05, * *p* ≤ 0.05, \*\**p* ≤ 0.01, *** *p* ≤ 0.001,**** *p* ≤ 0.0001).

ARi treatment of co-cultures of PCa cells significantly increased immune-mediated PCa killing at 48 hr, independent of PCa cell line AR status and sensitivity to ARi (**Fig.1 A-D**). Enzalutamide significantly increased NK cell-mediated killing of LNCaP at 48 hours (cell death rate: 15.6% ± 3.9 (control[C]) vs 48.3% ± 8.3 (enza); p<0.0001). This effect was also observed in ARi-resistant cell lines PC3 (3.8% ± 1.27 (C) vs. 11.05±4.6 (enza); p<0.0017), 22Rv1 (17.07% ± 3.35 (C) vs 52.67 ± 8.96 (enza); p<0.0001), and DU145 (13.45% ± 2.37 (C) vs 34.8 ± 4.92 (enza); p<0.001).

### The ARi immune enhancement and killing of PCa cell lines by NK cells is dependent on IFN-γ and TRAIL

The NK-92 cell line was treated with 10 nM of DHT for 72 h before enzalutamide or darolutamide, The culture supernatant was collected 24h after the addition of enzalutamide or darolutamide to measure cytokine concentrations. The secretion profile was assessed in monoculture conditions without the PCa cell lines (**Fig. 2A**).

**Figure 2.**
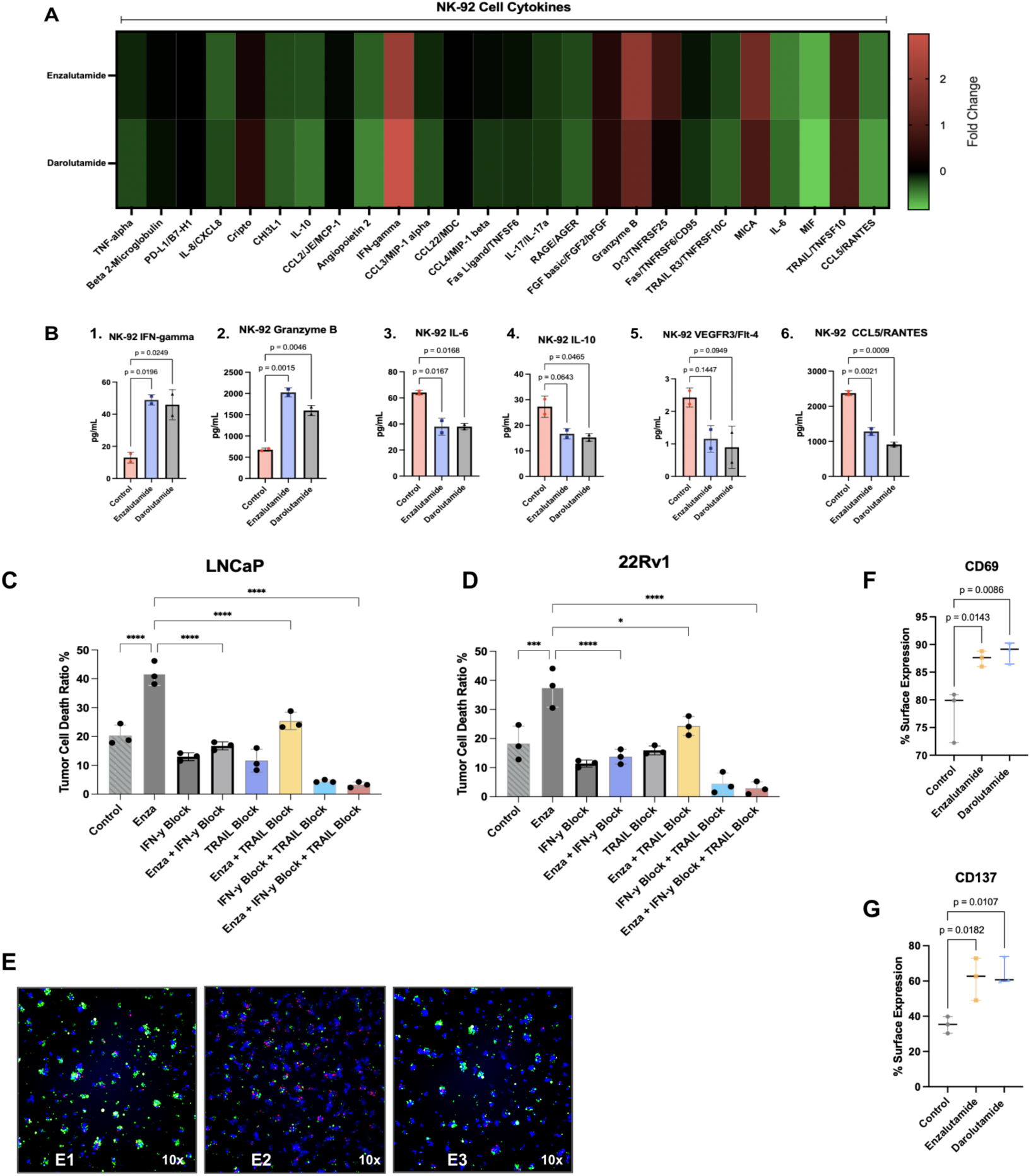
The ARi immune enhancement killing of PCa cell lines by NK-92 cells is mediated by IFN-γ and TRAIL. A: Fold change heat map displaying cytokine changes from the NK-92 cell line treated with enza and daro over 24h. B: Enza and daro significantly increase the secretion of cytotoxic cytokines from NK-92 cells (B1, B2). Enza and daro also reduced the secretion of immunosuppressive cytokines from the NK-92 cell lines (B4, B5). B6: Enza and daro reduced NK-92 cell line secretion of the T cell chemokine CCL5. C and D: IFN-γ blockade reduced the enza enhancement of NK-mediated immune killing of PCa. The TRAIL blocking mAb (RIK-2) partially reduced the enza immune enhancement effect. The dual blockade of TRAIL and IFN-γ reduces considerably the NK cell cytotoxic killing of PCa cell lines. E: Representative images of 22Rv1 + NK-92 (E1), 22Rv1 + NK-92 + Enza (E2), and 22Rv1 + NK-92 + Enza + IFN-γ block (E3). F-G: ARi increase the expression of NK cell activation markers, CD69 and CD137 (48h). (ns p>0.05, * p≤ 0.05, **p ≤ 0.01, p ≤ 0.001, **** p ≤ 0.0001)

Darolutamide and enzalutamide increased NK-92 cell secretion of IFN-γ ([C]: 12.98 pg/ml ± 3.4; Enza: 48.88 pg/ml ± 3.18; Daro: 45.6 pg/ml ± 9.4) and granzyme B ([C]: 676.6 pg/ml ± 36.2; Enza: 1,027 pg/ml ± 105.5; Daro: 1,599 pg/ml ± 118.6), and decreased the secretion of the immunosuppressive cytokine IL-10 ([C]: 27.3 pg/ml ± 4.16; Enza: 16.6 pg/ml ± 2.01; Daro: 15.24 pg/ml ± 1.48) (**Fig. 2B 1-4**). Decreases in VEGFR3 ([C]: 2.427 pg/ml ± 0.2913; Enza: 1,153 pg/ml ± 0.4098; Daro: 0.8933p g/ml ± 0.6510) were not significant. There was a significant decrease in CCL5 ([C]: 2,373 pg/ml ± 70.7; Enza: 1,281 pg/ml ± 110.7; Daro: 912 pg/ml ± 66.10) (**Fig. 2B 5,6**). These results support the role of ARi in activating NK cells and enhancing their cytotoxic function through the secretion of pro-inflammatory cytokines including IFN-γ.

To investigate the role of IFN-γ in mediating this ARi-induced enhancement of NK cell-mediated PCa cell killing, blocking IFN-γ monoclonal antibody (mAb; 10 µg/ml) was combined with ARi in co-culture experiments. Doses of blocking IFN-γ mAb were based on previous experiments and co-culture assays [18–20]. IFN-γ blockade reversed the immune enhancement effect of enzalutamide (**Fig. 2C, D**) in LNCap and 22Rv1 cells.

Based on the known role of NK cells in secreting TNF-related apoptosis inducing ligand (TRAIL) [21, 22] and the observed upregulation of the soluble form of a TRAIL receptor TRAIL-R2 by ARi on our cytokine analysis (**Fig. 3B**), we investigated the dual blockade of IFN-γ and TRAIL with RIK-2 (TRAIL-CD253 neutralizing mAb). Co-culture of PCa and NK cells in the presence of enzalutamide, IFN-γ mAb, and RIK-2 (TRAIL inhibitor) reduced the NK cell-mediated killing of PCa cells (**Fig. 2C, D**). In the presence of enzalutamide, the blockade of TRAIL diminished immune killing, but to a lesser degree than IFN-γ mAb blockade alone (**Fig. 2C, D**). The dual blockade, targeting TRAIL and IFN-γ, diminished NK cell PC cell killing almost entirely, even in the presence of ARi (**Fig. 2C, D**).

**Figure 3.**
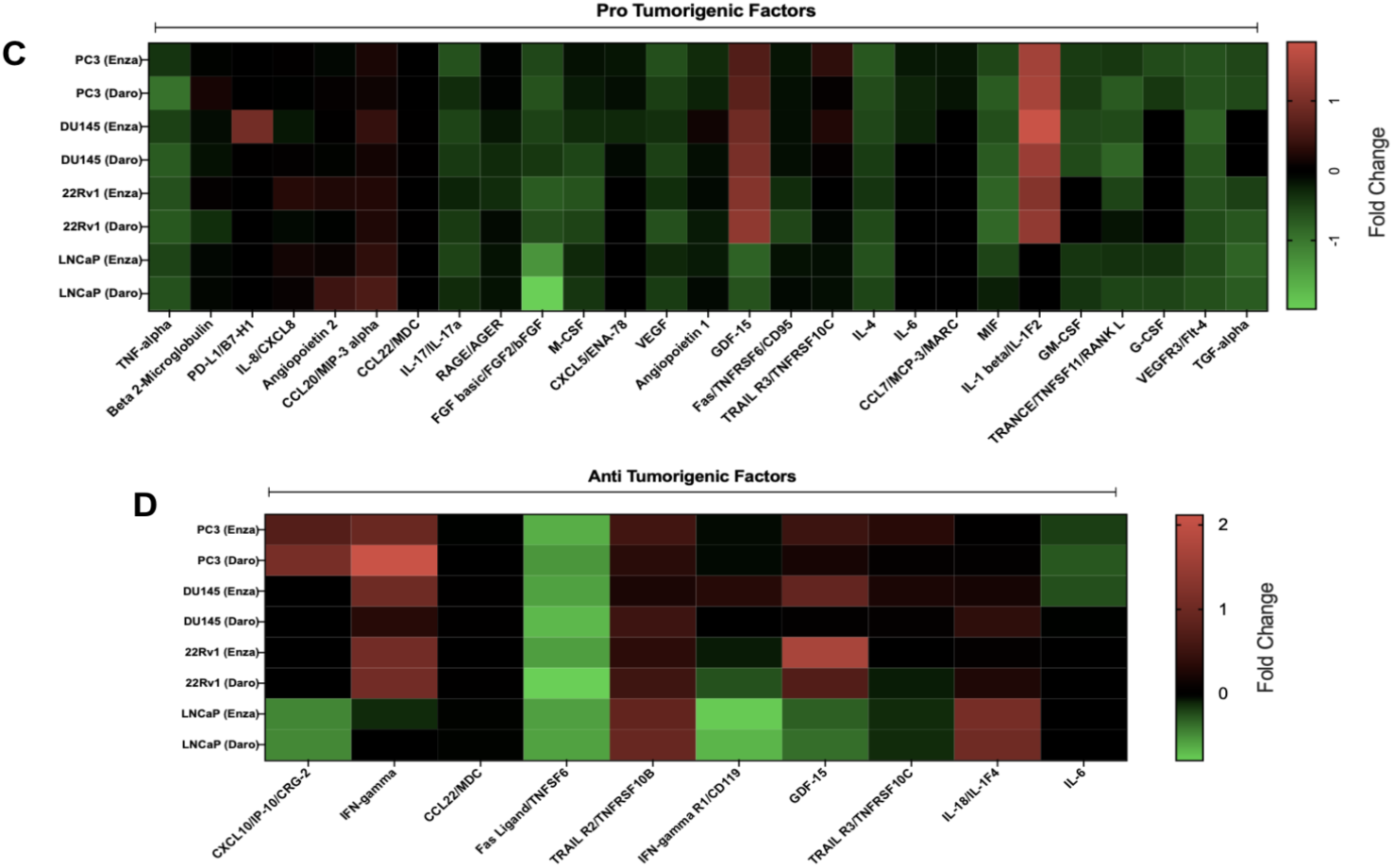
ARi modulates apoptotic and antiapoptotic proteins as well as pro and antitumorigenic cytokines in PCa cell lines. C and D: PCa cell lines (PC3, DU145, 22Rv1, and LNCaP) were treated with ARi for 24 h, and cytokines from the supernatant were analyzed and displayed as a heat map. These tumor-secreting cytokines were collected without the presence of NK cells. Absolute cytokine changes from the PCa cell lines are depicted in Supp. Fig. 3.

We also examined surface expression of the NK cell activation markers CD69 and CD137 following the treatment of NK cell monocultures with ARi (**Fig. 2F, G**). Enzalutamide and darolutamide increased the expression of CD69 and CD137 (CD69 [C]: 77.7% ± 4.75, enza: 87.4% ± 1.4, daro: 88.63% ± 1.94, p=0.009; CD137 [C]: 35.1% ± 4.77, enza: 61.54% ± 12, daro: 64.87% ± 7.96, p=0.01).

### Androgen receptor inhibitors modulate pro- and anti-tumorigenic cytokines in PCa cell lines

We investigated the effects of ARi on PCa cells cytokine secretion to assess a possible priming effect of treatment, which could sensitize these cells to NK cell killing. Each cell line was treated with 10 nM of DHT for at least 72 hours before treatment with enzalutamide or darolutamide. The results are depicted in **Fig. 3C, D** and **Supp. Fig 2**. Treatment of PCa cell lines with enzalutamide and darolutamide for 24h increased the secretion of immune-activating cytokines such as IL-2 and reduced the secretion of pro-tumorigenic and immunosuppressive cytokines such as VEGF, M-CSF, IL-8, CXCL10 and GDF-15 (**Fig. 3**; **Supp. Fig. 2A-D**). Additionally, DU145, 22Rv1, and PC3 cell lines had upregulation of IFN-γ. ARi downregulated soluble FasL and increased TRAIL-R2 and R3. Upregulation of GDF-15 by ARi was limited to ARi-resistant prostate cells (PC3, DU145, and 22Rv1). GDF-15 is implicated in the immunosuppressive prostate cancer TME and the progression of benign prostate hyperplasia to adenocarcinoma [18–20, 23, 24].

### NK cell activation is enhanced post-ARi treatment in patient samples

To investigate the effects of androgen deprivation therapy (ADT) on the activation of peripheral blood NK cells, blood samples were collected from patients diagnosed with castration-sensitive metastatic prostate cancer before initiation of ADT and approximately 26 days after patients had achieved castration levels of testosterone (median time between sample collections: 26.3±8.7 days). Patient characteristics are summarized in **Fig. 4A**. Per standard of care, ADT was implemented with the administration of luteinizing hormone-releasing hormone (LHRH) agonists (e.g., leuprolide) or antagonists (e.g., degarelix). Patient-derived NK cells were analyzed by flow cytometry. After ADT, the patients displayed a greater number of granzyme B+NK cells (pre-ADT: 20.53% ± 2.77; post-ADT: 37.24% ± 6.9; p=0.0017), and perforin+ NK cells (pre-ADT: 4.26% ± 0.92; post-ADT: 54.47% ± 6.47; p<0.001). The double-positive NK cell population (positive for granzyme B and perforin) also increased after ADT (pre-ADT: 3.54% ± 0.42; post-ADT: 38.15% ± 7.96; p=0.001) (**Fig. 4C, D**).

**Figure 4.**
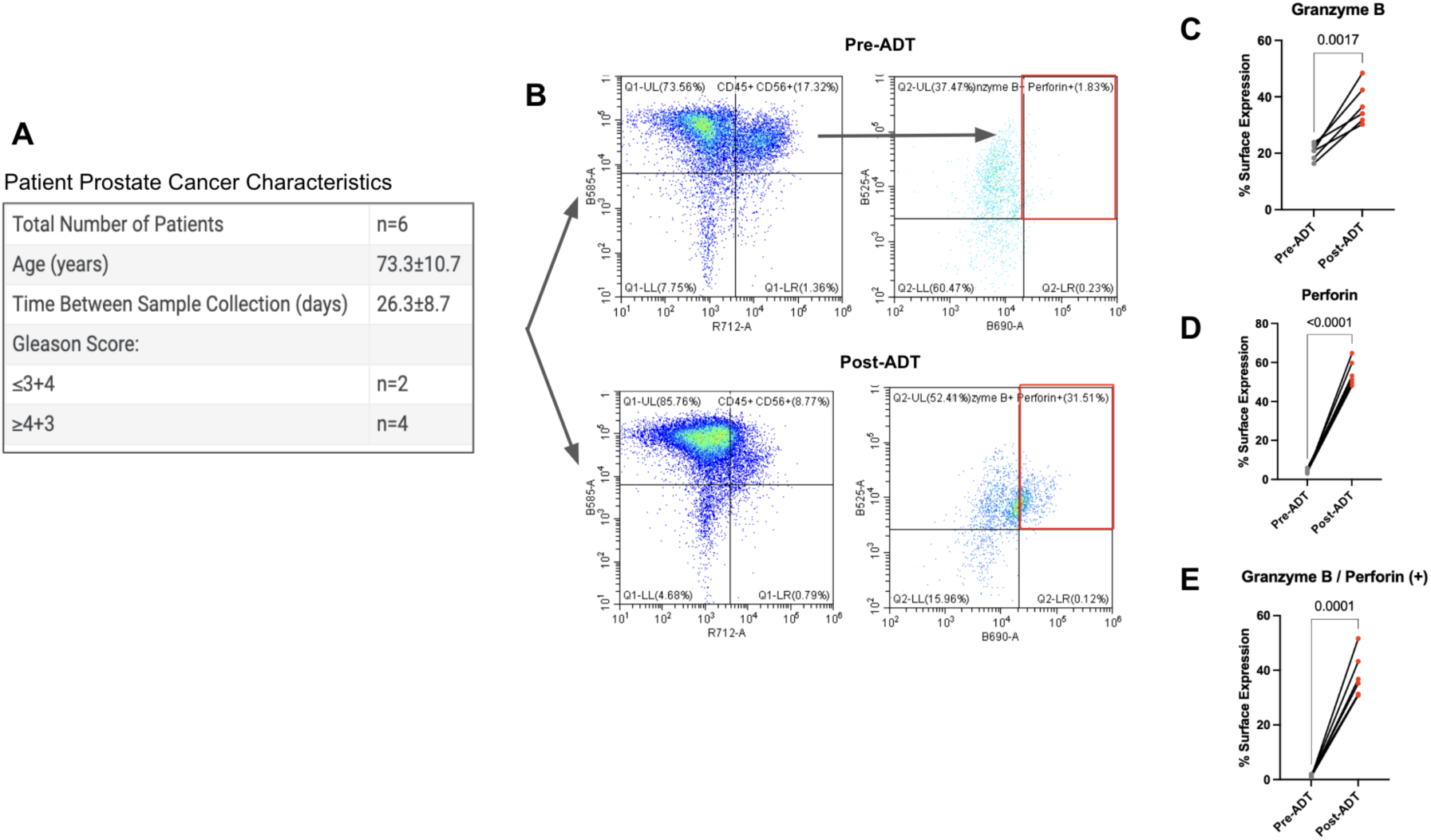
Androgen deprivation therapy (ADT) increases patient-derived NK cell cytotoxic markers. (A) Patient characteristics. Metastatic castration-sensitive prostate cancer patients (n=6) had whole PBMCs and NK cell isolation performed before and after the start of ADT. The activation profile of these NK cells was evaluated by flow cytometry. NK Cells were gated on CD56+CD45 live PBMC population, and a representative matched patient is depicted in B. (C) Granzyme B, (D) Perforin, and double positive population (E) displayed increased expression after ADT. A paired t-test was performed, and the results indicate the matched patient cohort (n=6) pre- and post-ADT.

### Blocking the NK cell receptor NKG2A with NKG2A blocking mAb monalizumab potentiates the ARi-induced immune-mediated killing of PCa cells

To evaluate a strategy to potentiate NK cell killing induced by ARi, we performed co-culture experimentsco-cultured NK cells with enzalutamide plus monalizumab, an NKG2A blocking mAb. Monalizumab significantly increased enzalutamide-induced immune enhancement irrespective of the ARi sensitivity of PCa cell lines. (**Fig. 5A, B**). Monalizumab as a single agent did not enhance NK cell killing (**Fig. 5A, B**) or impact the viability of PCa or NK cells (data not shown).

**Figure 5.**
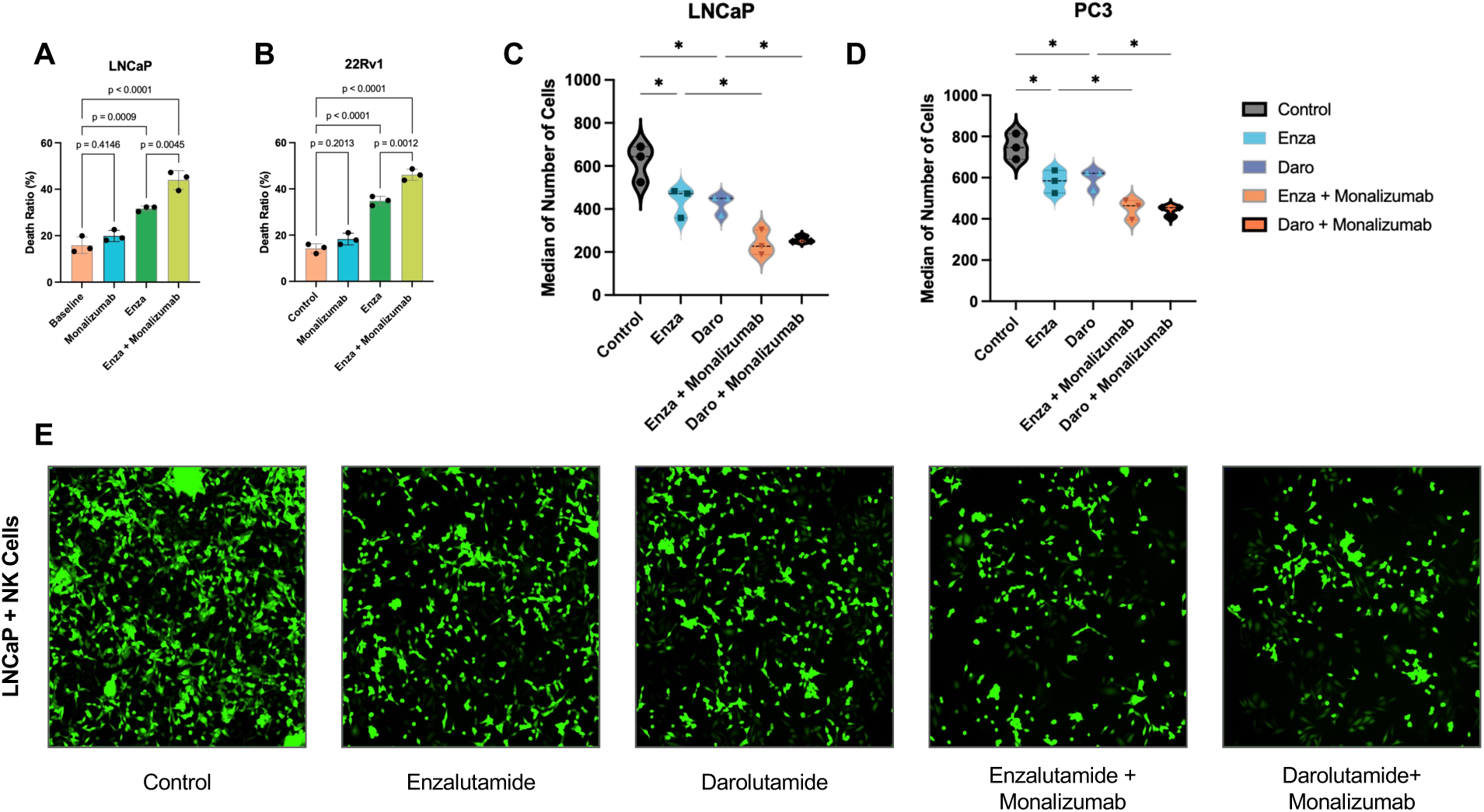
Blocking the NK cell receptor NKG2A with monalizumab potentiates the ARi-induced NK cell activation and killing of PCa cells. PCa cell lines (A: LNCaP and B: 22Rv1) were treated with enzalutamide (10 mM) and monalizumab (15 µg/mL) in the presence of NK-92 cells (1:1) ratio. Monotherapy with monalizumab did not significantly enhance the NK cell killing of PCa cell lines (A, B). ARi and monalizumab significantly enhanced the NK cell-mediated killing of PCa cell lines. GFP-expressing PCa cells were plated with patient-derived NK cells (1:1) ratio. Enza and darolutamide enhanced the killing of the PCa cell lines over 24 h. The addition of monalizumab in combination with enza and daro potentiated the killing of PCa (C, D). E. Representative images of the GFP-expressing LNCaP cell line co-cultured with patient-derived NK cells and treated with ARi and monalizumab (Anti-NKG2A). (ns p>0.05, * p≤ 0.05, **p ≤ 0.01, p ≤ 0.001, **** p ≤ 0.0001).

To validate an additive effect of ARi and monalizumab enhancing NK cell cytotoxicity, we performed a co-culture assay with co-cultured PCa cell lines (including ARi-sensitive LNCaP and ARi-resistant PC3) and patient-derived NK cells. The NK cells were negatively selected from whole blood from patients with metastatic prostate cancer prior to initiation of androgen deprivation therapy. The addition of enzalutamide or darolutamide enhanced the cytotoxicity of NK cells, and the combination of ARi with monalizumab potentiated immune killing (**Fig. 5C-E**). Even though this effect was evident irrespective of the AR status of the cell line, the AR-sensitive cell line, LNCaP, displayed greater sensitivity to the addition of monalizumab in combination with enzalutamide or darolutamide (**Fig. 5C-E**).

### Androgen receptor blockade upregulates HLA-E in androgen-signaling sensitive, but not in androgen-independent, cell lines

The effect of ARi on the expression of NKG2A ligand, HLA-E, was investigated by flow cytometry on PCa cells treated with enza and daro. ARi treatment increased the surface expression of HLA-E in LNCaP ([C]: 28.57 ± 2.4×10^3^, enza: 37.38 ± 1.7×10^3^ and daro: 39.47 ± 3.39×10^3^, p=0.004) (**Fig. 6A**), and did not affect HLA-E expression of ARi-resistant cell line 22Rv1 ([C]: 34.76 ± 5.1×10^3^, enza: 41.14 ± 7.6×10^3^ and daro: 30.5 ± 2.69×10^3^, p=0.14) (**Fig. 6B**).

**Figure 6.**
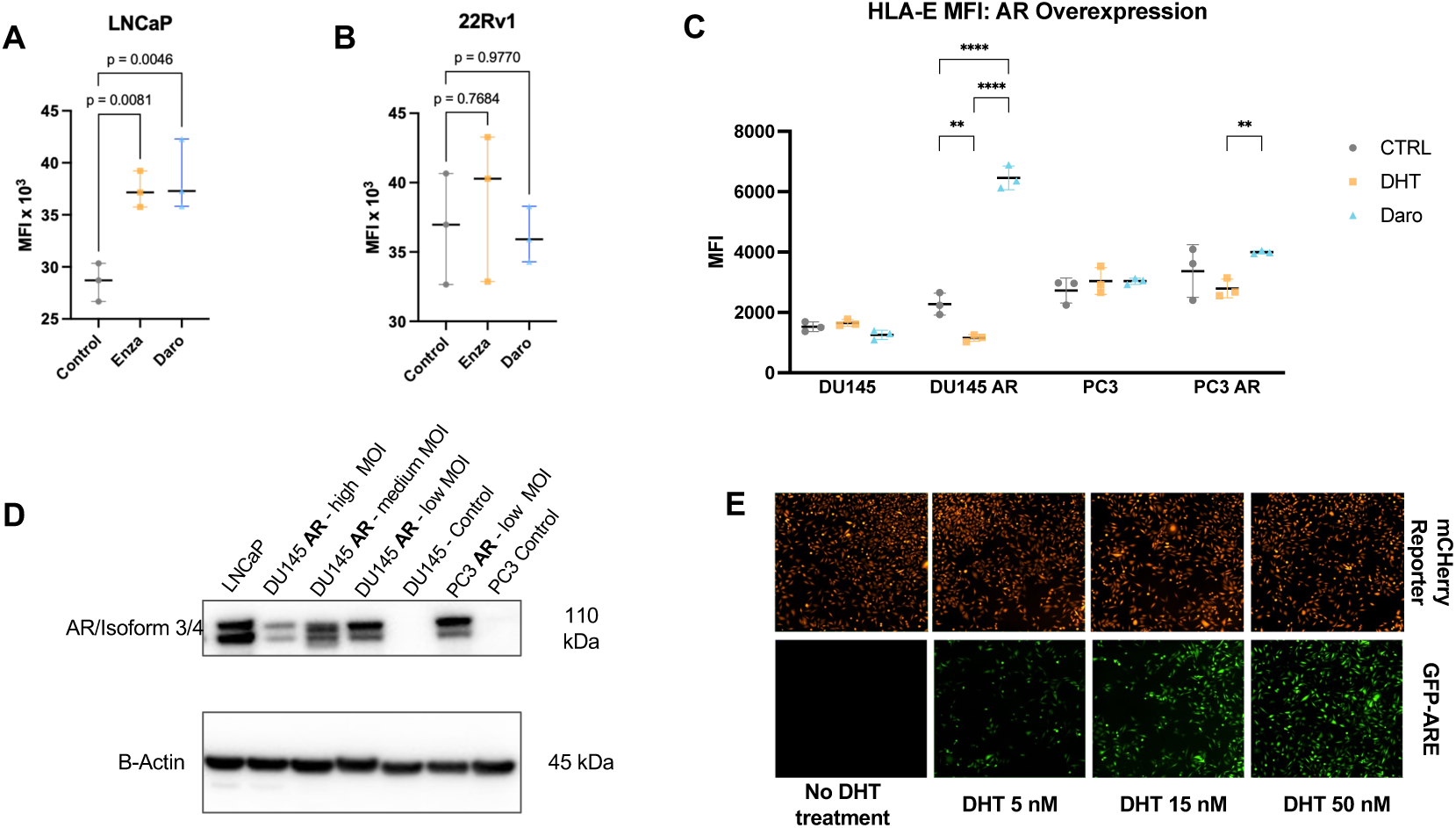
Prostate cancer cells’ expression of HLA-E, the ligand of NKG2A, is modulated by androgen signaling. HLA-E mean fluorescence intensity (MFI) on LNCaP (A) and 22Rv1 (B) cells treated with enza (15uM) and daro (20 uM) for 48 h. To evaluate the AR signaling dependence on the surface expression of HLA-E, the AR negative cell lines (PC3 and DU145) were transduced with AR (D) and an AR activity reporter (E). Low MOI cells were used for downstream experiments. The m-Cherry AR transduced reporter expressed on PC3 (E) and DU145 cell lines was functionally assessed and exhibited a GFP signal when cells were treated with increasing DHT doses. C: The AR-negative cell lines PC3 and DU145 and the new AR-responsive cell lines PC3-AR cell line were evaluated for surface expression of HLA-E upon DHT treatment and ARi with darolutamide. The AR-negative DU145 cell line did not express significant changes in HLA-E surface expression (C), but the AR-responsive isogenic model (DU145-AR) displayed increased HLA-E upon ARi compared to DHT-treated. Similar findings were also observed on the PC3 (AR-responsive) cell lines (C). Statistical significance in (C) was determined by two-way ANOVA with Tukey’s correction for multiple comparisons (**p<0.01, ****p<0.0001).

To evaluate the modulation of the androgen receptor and tumor HLA-E expression, we stably transduced AR negative cell lines PC3 and DU145 with a mCherry-AR reporter driven by a GFP-Androgen Response Element (ARE). After stably transducing these reporter-containing cells with an AR receptor (**Fig. 6D**), the function of the ectopic AR was validated by treating the AR+-constructed cell lines with DHT at different concentrations over 72h. The expression and targeting of ARE were confirmed by increased expression of GFP+ cells (**Fig. 6E**).

Using AR+ constructed PC3 and DU145, we sought to determine the dependence of HLA-E expression and possible modulation by AR. As seen previously with the ARi resistant 22Rv1 cell line, ARi did not increase the HLA-E expression of non-AR-transduced PC3 and DU145 cell lines (**Fig 6C**). In the AR-transduced DU145 line, however, HLA-E expression increased upon ARi treatment, like the AR-sensitive LNCaP cell line (**Fig. 6**). A similar, but less pronounced, trend was observed in AR-transduced PC3, which demonstrated a statistically significant increase in HLA-E between DHT and daro-treated conditions. These results suggest that AR antagonism can increase the expression of HLA-E in PCa cells depended on androgen signaling. Higher expression of HLA-E in turn inhibits the NK cell cytotoxicity via NKG2A receptors favoring immune scape of prostate cancer tumors. This finding complements the earlier result that ARi in combination with NKG2A blockade yielded a better killing profile in the presence of NK cells in AR-dependent than in AR-insensitive prostate cancer cells. Increased HLA-E in PCa cells mediated by ARi could represent a novel mechanism of resistance or a contributor to the immunosuppressive “cold” phenotype of prostate cancer TME.

To explore a potential epigenetic regulation of HLA-E expression by androgen signaling, the HLA-E expression was evaluated on the PC3 and LNCaP after DHT treatment with and without pan-HDAC inhibitors vorinostat and panobinostat. Experiments were conducted with noncytotoxic doses of vorinostat (0.3 μM) and panobinostat (3 nM; **Supp. Fig. 3**). Both cell lines were kept in culture for at least five days in the presence of 30 nM of DHT before the experiments (**Fig. 7**). DHT alone or combined with HDACi did not change the HLA-E expression of AR negative cell line PC3 (**Fig. 7A**). However, DHT reduced significantly the HLA-E expression on the AR responsive cell line LNCaP and this effect was blocked when DHT was combined with HDACi (**Fig. 7B**). These results suggest that HLA-E regulation in the ARi-sensitive cell lines may depend on epigenetic regulation by HDACs.

**Figure 7.**
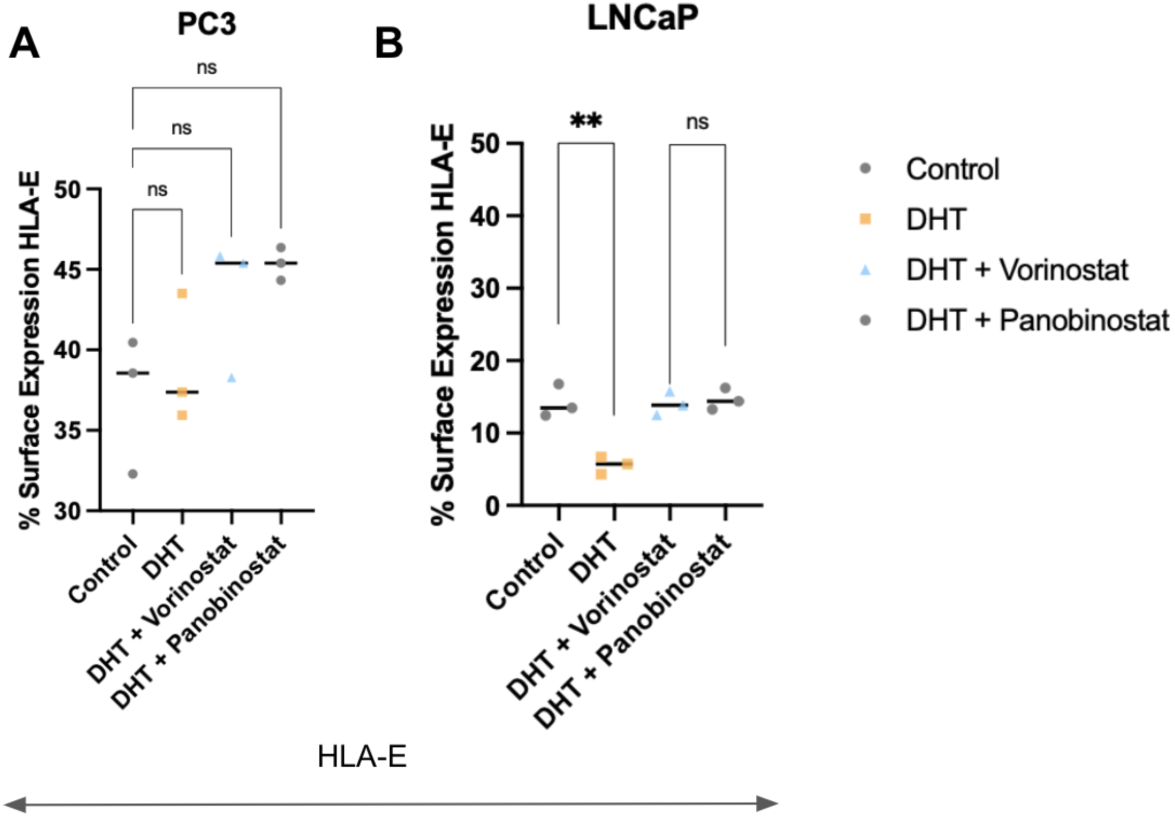
HDAC inhibitors block androgen-mediated repression of HLA-E in LNCaP cells. The PC3 and LNCaP cell lines were kept in DHT (30 nM) supplemented media for at least five days before experiments. (A-B) Surface expression of HLA-E (48h) was evaluated on the AR-responsive cell lines LNCaP and the AR-non-responsive PC3 cell line after HDACi (vorinostat 0.3 μM; panobinostat 3 nM). HLA-E was also assessed after DHT treatment with and without HDACi. (ns p>0.05, * p≤ 0.05, **p ≤ 0.01, p ≤ 0.001, **** p ≤ 0.0001).

### HLA-E ablation enhances NK cell response to androgen receptor inhibitors *in vivo*

To study the anti-tumor effect of disrupting NKG2A-HLA-E signaling in combination with ARi, we conducted an *in vivo* study using a syngeneic prostate cancer mouse model with the RM-1 cell line (**Fig. 8A**). We genetically ablated the murine HLA-E locus, H2-T23, using CRISPR-Cas9 and validated a reduction in expression of its protein product, QA-1b, by western blot (**Fig. 8B**). Following the establishment of sgRNA control and HLA-E knockout (KO) RM-1 tumors, mice were chemically castrated with degarelix and treated with darolutamide or vehicle control for 6 days (**Fig. 8A**). At day 6, HLA-E KO tumors exhibited reduced growth compared to wild-type tumors, suggesting a potential impact on the innate immune activation. As expected, darolutamide significantly reduced the growth of HLA-E wild-type tumors. Notably, the tumor growth inhibition of darolutamide was further enhanced in HLA-E KO tumors compared with wild-type tumors. These results are consistent with the additive effect of monalizumab in the darolutamide-mediated NK cell killing. Interestingly, darolutamide increased NK cell infiltration, as measured by NK1.1 expression, in wild-type tumors but not HLA-E KO tumors (**Fig. 8D**). In contrast, darolutamide significantly increased granzyme B expression in HLA-E KO tumors compared to wild-type tumors treated with darolutamide (**Fig. 8E**). The dissociation between NK1.1 and granzyme B suggests that HLA-E loss primarily enhances NK cell cytotoxic activation rather than NK cell recruitment. Given the canonical inhibitory role of HLA-E-NKG2A pathway in NK cells, the increased granzyme B in HLA-E KO tumors likely reflects enhanced NK cell effector function rather than CD8 T cell activation. These findings suggest that ARi promotes NK cell infiltration, whereas HLA-E loss preferentially enhances NK cell cytotoxic function consistent with its role as an inhibitory checkpoint ligand, resulting in an additive increase in NK cell activation as measured by granzyme B staining.

**Figure 8:**
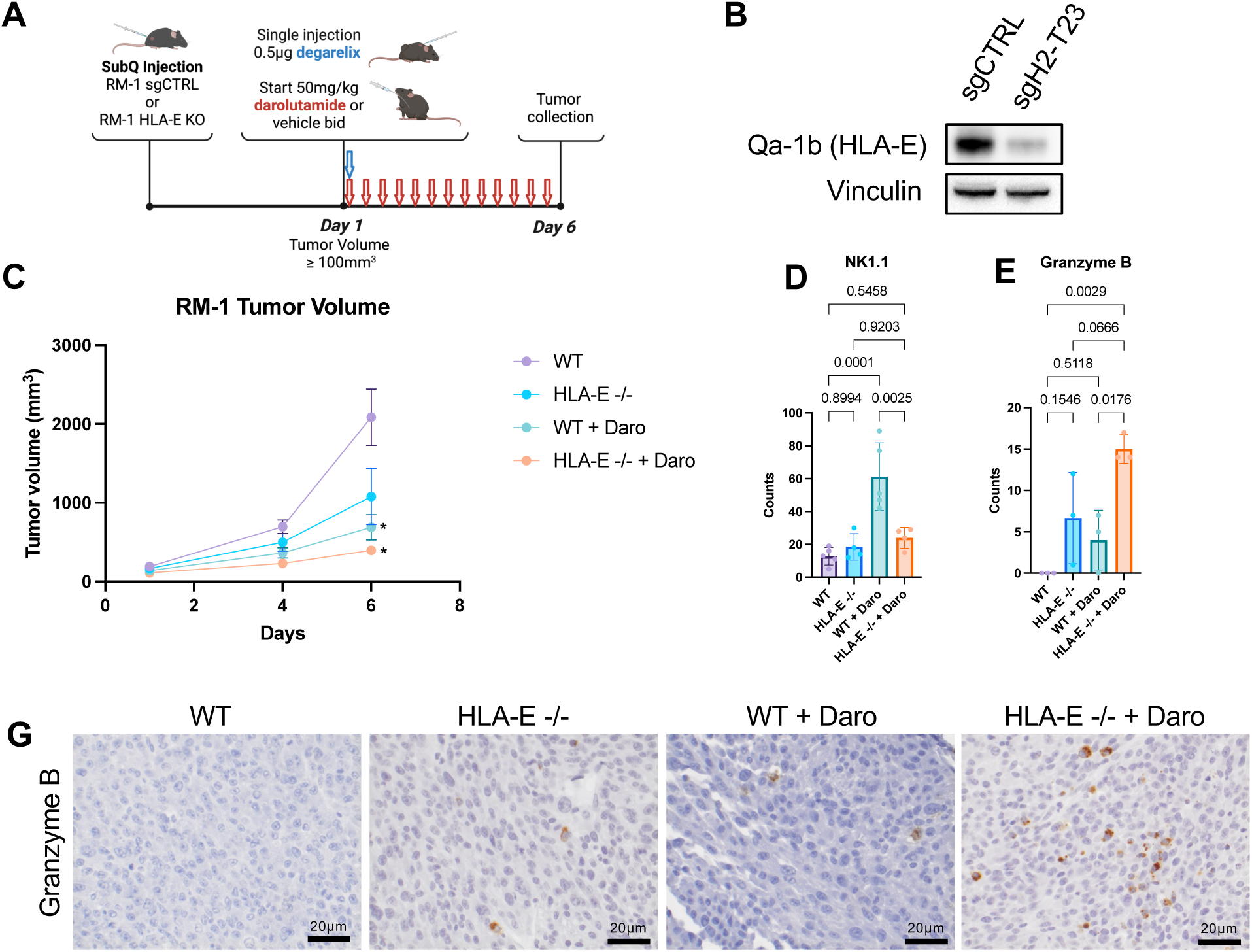
Darolutamide synergizes with HLA-E ablation in vivo. A: Schematic of mouse study. B: Western blot analysis of RM-1 mouse prostate cancer cells following transfection of Cas9 ribonucleoprotein complexed with HLA-E targeting sgRNA or a nontargeting control. C: Subcutaneous RM-1 tumor volumes collected at days 1, 4, and 6 during the study, where day 1 is the first day of darolutamide treatment. D: Quantification of (D) NK1.1 and (E) Granzyme B staining in HLA-E wild-type or knockout RM-1 cells treated with darolutamide or vehicle for 6 days, and (G) corresponding representative images.

## Discussion

Our results demonstrate that androgen signaling blockade activates NK cells in vitro and enhances the killing of PCa cells in co-culture experiments. The AR inhibitors enzalutamide and darolutamide increased NK cell secretion of cytokines IFN-γ and TRAIL that mediated PCa killing. The blockade of IFN-γ inhibited this effect significantly, and the dual blockade of IFN-γ and TRAIL had further additive effects. Monalizumab, a monoclonal antibody blocking the immune inhibitory receptor NKG2A on NK cells, potentiated the NK cell cytotoxicity induced by ARi. The AR inhibitors also modulated PCa cells by decreasing the secretion of pro-tumorigenic factors (e.g. IL-4, TNF-a, VEGF, MIF, M-CSF, and GM-CSF). Our results also reveal a novel androgen signaling regulation of HLA-E expression in PCa cells. HLA-E is a nonclassical HLA class I molecule and the main ligand of the NKG2A receptor [25]. HLA-E is frequently overexpressed in tumors and contributes to immune escape in the TME by inhibiting NK cells and NKG2A-expressing CD8 T cells [26]. ARi-induced HLA-E expression in prostate cancer cells can protect them from NK lysis, contribute to a “cold” immune environment, and may represent a mechanism of resistance enabling the persistence of prostate cancer cells during treatment with AR inhibitors as well as ADT.

Our findings elucidate the immune-modulatory role of ARi therapy on NK cells, complementing results describing the effect of ADT and AR blockade on T cell function. Building on a clinical trial investigating the combination of anti-PD1 (pembrolizumab) with enzalutamide for the treatment of mCRPC, Guan and colleagues showed that enzalutamide significantly enhanced the anti-tumor effect of anti-PD-L1 plus ADT in animal models. These effects were mediated by a direct effect of enzalutamide increasing T cell secretion of IFN-γ and granzyme B. The group demonstrated that the androgen receptor interacts with IFN-y and granzyme B genes in open chromatin regions (OCRs) of memory CD8 T cells, blocking the rapid production of IFN-γ and granzyme B enabled by these OCRs upon TCR stimulation. Blocking the androgen-mediated suppression of IFN-y production in T cells with enzalutamide improved the anti-tumor response elicited by PD-L1 inhibitor, helping overcome the relative resistance of prostate cancer to checkpoint inhibitors *in vivo* [8]. Other results corroborate the androgen immune suppressive effects on T cells. ADT increased tumor infiltration of IFN-γ-expressing T cells in the prostate TME. Enzalutamide activates IFN-γ signaling pathways and decreases the frequency of immunosuppressive cells in peripheral blood in mononuclear cells isolated from patients with mCRPC [27, 28]. Our results show a direct effect of AR inhibitors increasing NK activation and secretion of IFN, which can also impact T cell function through the functional crosstalk between these two cell lines, enabling an effective anti-tumor response [29]. IFN-γ produced by NK cells promotes an anti-tumor T helper cell type 1 (Th1) polarization in CD4+ T cells [30]. Activated NK cells stimulate CD4+ T cell proliferation by interacting with OX40/OX40L [31] and indirectly influence T cell response by regulating differentiation of dendritic cells. NK cells can induce a type 17 polarization in CD8+ T cells, characterized by the ability to produce IFN-γ and IL-17A through priming of dendritic cells [32]. These findings suggest that therapeutic strategies targeting the activation of CD8 T cells and NK cells could overcome the immune suppressive TME in prostate cancer and lead to meaningful anti-tumor effects. One such strategy could be the combination of AR inhibitors plus monalizumab and anti-PD1/PD-L1 agents. Monalizumab is in clinical development for the treatment of lung cancer and other solid tumors in combination with durvalumab (anti-PD-L1) as a novel strategy to harness the innate immune response and activate NK cells, leading to clinically meaningful tumor response and immune infiltration [13, 33–35].

The importance of NK function in PCa and the therapeutic potential of targeting these cells is highlighted by results showing their impact on the clinical outcomes of patients. In a recent pan-cancer analysis, natural killer cell infiltration in PCa tumor specimens was associated with improved OS (HR 0.46, 95% CI 0.38–0.56, p=0.0001) [36]. The study also found that NK cell infiltration was associated with a 1.4-2.1–fold increased expression of immunomodulatory receptors LAG3 and TIGIT (p=0.0001). Another analysis of a large cohort of tumor specimens of patients with mCRPC showed that tumor infiltration by cytotoxic NK cells and higher expression of activating receptors NKp30 and NKp46 in NK cells isolated from the peripheral blood was associated with improved overall survival and a longer interval to development of castration resistance [37]. Cytokine signaling in the TME, especially IL-6, was associated with resistance to NK cell-mediated cytotoxicity via modulating PD-L1 and NKG2D ligand levels in PCa cells [38]. Peripheral blood NK cell dysfunction was also associated with poor clinical outcomes of PCa and higher disease stage [39]. These findings suggest that modulation of NK cell activation could help overcome and challenge the limited benefit of T cell-centric strategies deployed by PD-1/PD-L1 or CTLA-4 inhibitors for treating mCRPC.

Novel strategies blocking the inhibitory receptor NKG2A expressed on NK and CD8+ cells are under clinical investigation to overcome resistance to anti-PD-1/PD-L1 checkpoint inhibition [40]. NKG2A is an inhibitory checkpoint expressed on the cell surface of nearly half of circulating NK cells, some activated CD8 T cells, and can be induced by cytokines such as IL-15 and IL-12 [41, 42]. It is a heterodimer, bound with CD94 in humans and mice, and recognizes the non-classical class I major histocompatibility complex (MHC-I) molecules of human leukocyte antigen (HLA)-E. HLA-E expression is present in normal tissue as a protective “self” signaling, and it is up-regulated in various malignancies, including prostate cancer [43–46]. The inhibitory function on NK cells is mediated through the binding of HLA-E to NKG2A/CD94, leading to the recruitment of the SHP-1 tyrosine phosphatase to the tyrosine-phosphorylated form of the intracytoplasmic tyrosine-based inhibitory motifs (ITIM). When phosphorylated, these motifs recruit phosphatases (SHP-1/2 or SHIP) responsible for transmitting inhibitory signals to immune effector cells. To our knowledge, we describe for the first time that HLA-E expression in prostate cancer cells is modulated by androgen signaling. AR inhibitors enzalutamide and darolutamide increased the expression of HLA-E in androgen-sensitive PCa cell lines. The ARi-driven upregulation of HLA-E could suppress NK cells and the innate immune response and contribute to the cold tumor microenviroment. Even though strategies enhancing NK cell cytotoxicity might display promising results, blocking the immune inhibitory HLA-E/NKG2A axis could result in additional benefits.

The limitations of our results include incomplete insight into the mechanisms of androgen receptor regulation of NK cell cytokine production. Whether this is a direct effect mediated by AR on ARE, a post-translational regulation, or is under epigenetic modulation also remains to be characterized. It would also be valuable to investigate the expression of HLA-E in human prostate tumor specimens pre- and post-androgen blockage to validate the observation in PCa cell lines. Further, given the diverse set of immune cells present in the TME, it would be valuable to profile the impact of Ari and HLA-E inhibition in additional immune cell types, such as CD4+ T cells and macrophages.

These results highlight the potential therapeutic implication of NK-activating strategies for the treatment of prostate cancer. The additive effect of androgen receptor inhibitors and monalizumab (anti-NKG2A) activating NK cells and enhancing PCa cell killing support further investigation of this combination in vivo with the potential for immediate clinical translation. Encouraging synergy of durvalumab and monalizumab in ongoing clinical trials for the treatment of lung cancer supports the clinical feasibility of this combination that could be expanded with the addition of androgen receptor inhibitors in advanced prostate cancer as a potential novel platform to modulate the cold TME in prostate cancer.

## Acknowledgements

W.S.E-D. is an American Cancer Society Research Professor and is supported by the Mencoff Family University Professorship at Brown University. This work was supported by an NIH grant (CA173453) to W.S.E-D. This work was presented in part at the 2023 meeting of the American Association for Cancer Research.

---

**Supplementary Figure 1.**
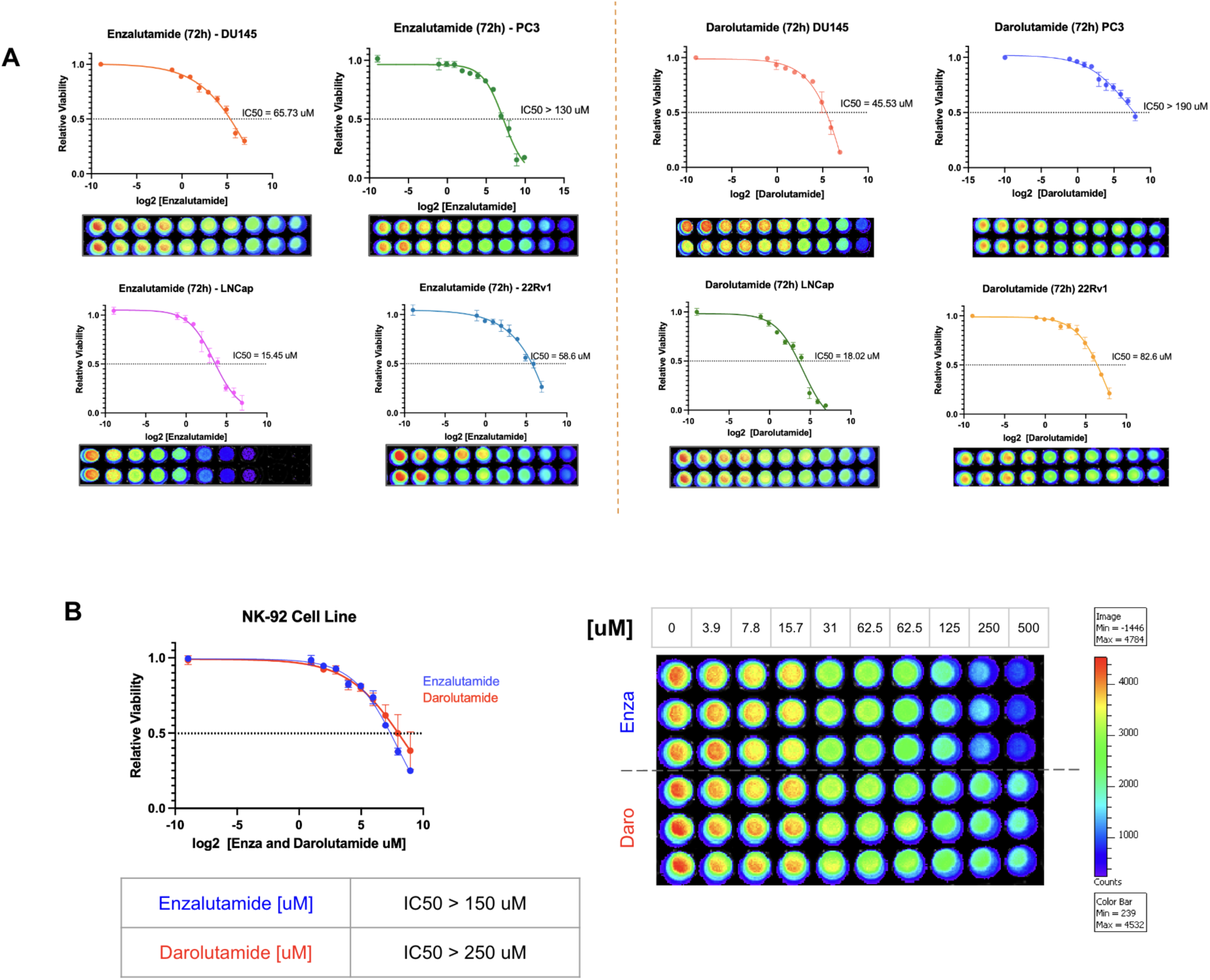
Enzalutamide and darolutamide viability assays with prostate cancer cell lines and NK-92 cell line. A - Cell Titer Glow (CTG) viability assay of PCa cell lines treated with enzalutamide or darolutamide. Five thousand cells were plated overnight and treated with increasing concentrations of enzalutamide and darolutamide after 24 h. Viability assay was performed at 72 h and normalized to the control. The IC50 was determined by nonlinear regression. B - NK-92 cell line viability was assessed upon enza and darolutamide treatment. Daro and enza displayed nontoxicity on NK-92 cells with doses used in the coculture assays (enza=10 μM and daro=20 μM).

**Supplementary Fig. 2.**
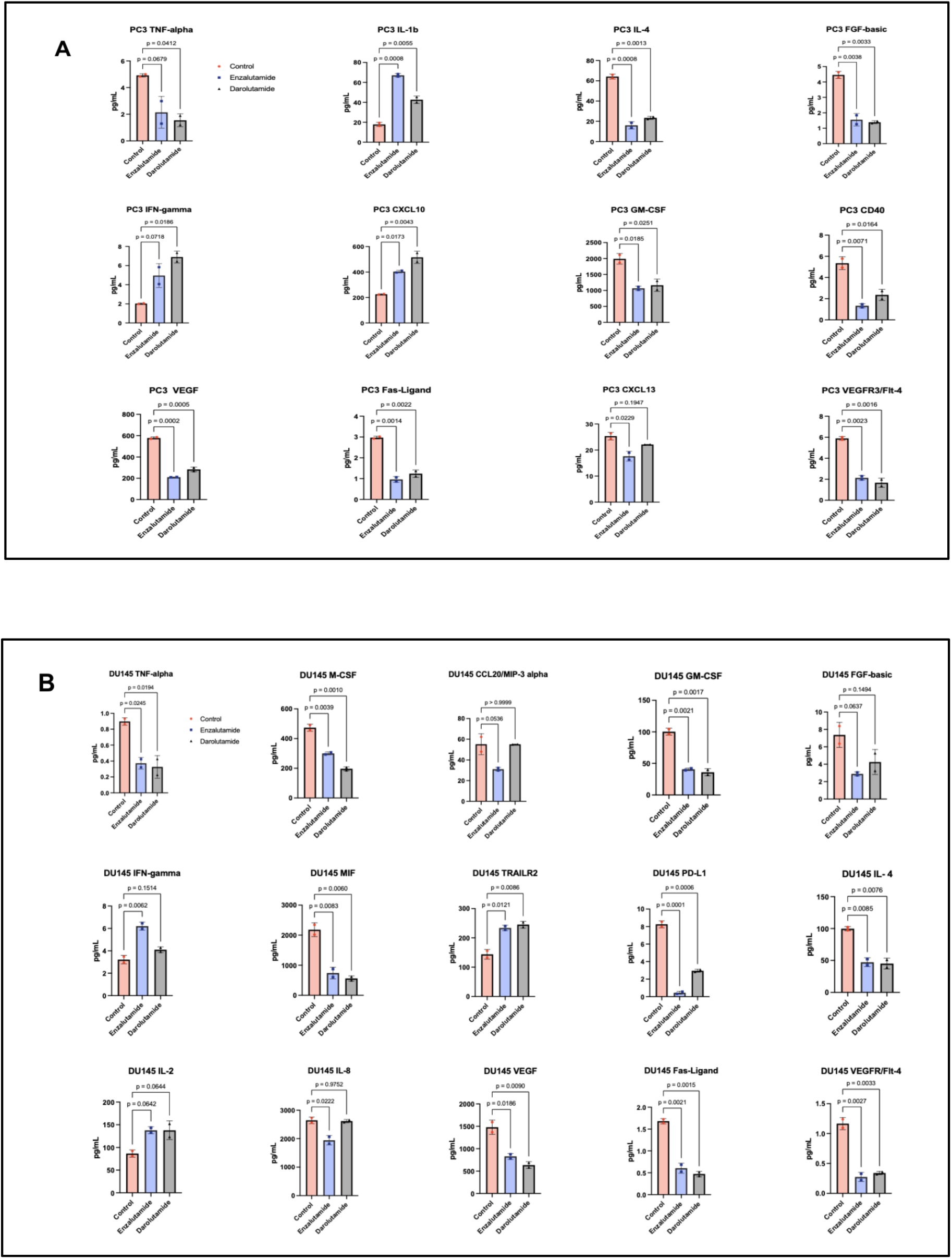
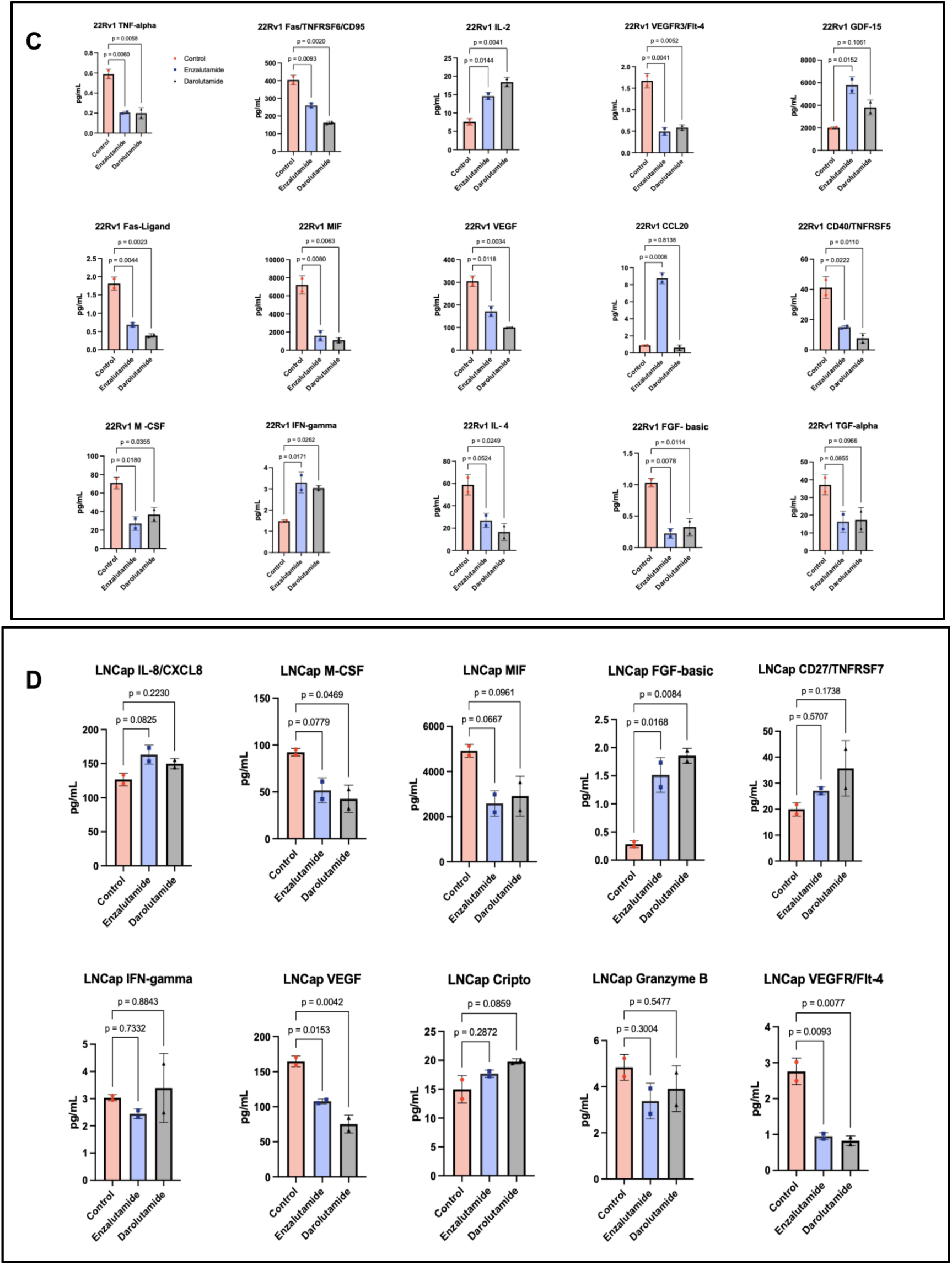
Cytokine secretion profile of PCa treated with Ari. Individual cytokine analysis from PCa treated with enza and darolutamide after 24 h. The PC3 (A), DU145 (B), 22Rv1 (C), and LNCaP (D) cell lines were treated with enza (10 μM) and daro (15 μM) for 24 h, and the supernatant was collected for cytokine profiling.

**Supplementary Fig. 3.**
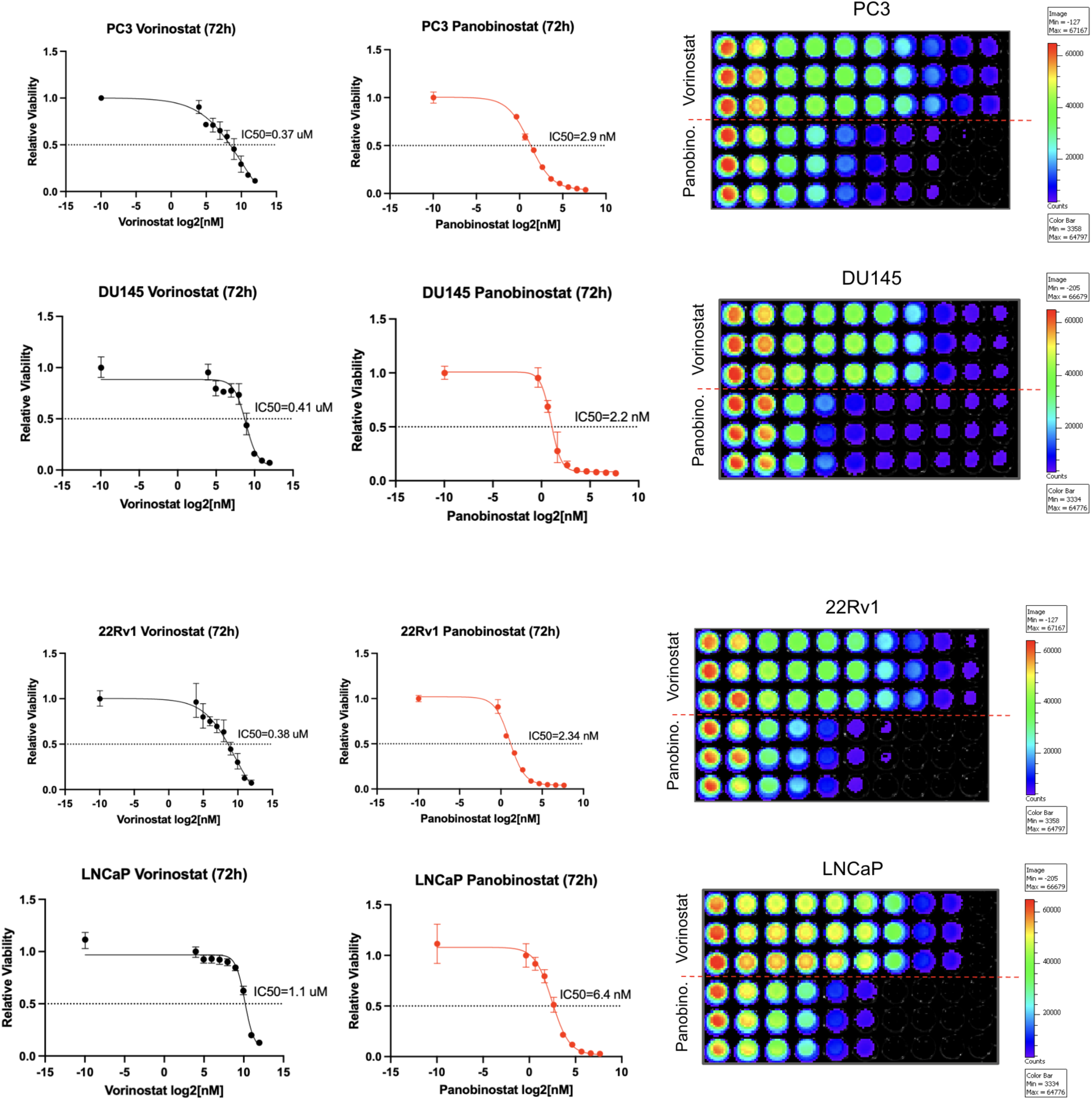
PCa cell lines viability with pan HDAC inhibitors (Panobinostat and Vorinostat), Cell Titer Glow (CTG) viability assay of PCa cell lines treated with panobinostat or vorinostat. Five thousand cells were plated overnight and treated with increasing concentrations of panobinostat or vorinostat after 24 h. Viability assay was performed at 72 h and normalized to the control. The IC50 was determined by non-linear regression.

## References

1. Siegel, R.L., et al., Cancer statistics, 2023. CA Cancer J Clin, 2023. 73(1): p. 17–48.

2. Freedland, S.J., et al., Real-world treatment patterns and overall survival among men with Metastatic Castration-Resistant Prostate Cancer (mCRPC) in the US Medicare population. Prostate Cancer Prostatic Dis, 2024. 27(2): p. 327–333.

3. Antonarakis, E.S., et al., Pembrolizumab for Treatment-Refractory Metastatic Castration-Resistant Prostate Cancer: Multicohort, Open-Label Phase II KEYNOTE-199 Study. J Clin Oncol, 2020. 38(5): p. 395–405.

4. Vitkin, N., et al., The Tumor Immune Contexture of Prostate Cancer. Front Immunol, 2019. 10: p. 603.

5. Karwacki, J., et al., Boosting the Immune Response-Combining Local and Immune Therapy for Prostate Cancer Treatment. Cells, 2022. 11(18).

6. Schweizer, M.T., et al., CDK12-Mutated Prostate Cancer: Clinical Outcomes With Standard Therapies and Immune Checkpoint Blockade. JCO Precis Oncol, 2020. 4: p. 382–392.

7. Barata, P., et al., Clinical activity of pembrolizumab in metastatic prostate cancer with microsatellite instability high (MSI-H) detected by circulating tumor DNA. J Immunother Cancer, 2020. 8(2).

8. Guan, X., et al., Androgen receptor activity in T cells limits checkpoint blockade efficacy. Nature, 2022. 606(7915): p. 791–796.

9. Kissick, H.T., et al., Androgens alter T-cell immunity by inhibiting T-helper 1 differentiation. Proc Natl Acad Sci U S A, 2014. 111(27): p. 9887–92.

10. Graff, J.N., et al., A phase II single-arm study of pembrolizumab with enzalutamide in men with metastatic castration-resistant prostate cancer progressing on enzalutamide alone. J Immunother Cancer, 2020. 8(2).

11. Henze, L., D. Schwinge, and C. Schramm, The Effects of Androgens on T Cells: Clues to Female Predominance in Autoimmune Liver Diseases? Front Immunol, 2020. 11: p. 1567.

12. Gubbels Bupp, M.R. and T.N. Jorgensen, Androgen-Induced Immunosuppression. Front Immunol, 2018. 9: p. 794.

13. Cho, M., et al., Durvalumab + monalizumab, mFOLFOX6, and bevacizumab in patients (pts) with metastatic microsatellite-stable colorectal cancer (MSS-CRC). Annals of Oncology, 2019. 30.

14. Cao, B., et al., Androgen receptor splice variants activating the full-length receptor in mediating resistance to androgen-directed therapy. Oncotarget, 2014. 5(6): p. 1646–56.

15. Kwilas, A.R., et al., Androgen deprivation therapy sensitizes triple negative breast cancer cells to immune-mediated lysis through androgen receptor independent modulation of osteoprotegerin. Oncotarget, 2016. 7(17): p. 23498–511.

16. Barton, V.N., et al., Multiple molecular subtypes of triple-negative breast cancer critically rely on androgen receptor and respond to enzalutamide in vivo. Mol Cancer Ther, 2015. 14(3): p. 769–78.

17. Mina, A., R. Yoder, and P. Sharma, Targeting the androgen receptor in triple-negative breast cancer: current perspectives. Onco Targets Ther, 2017. 10: p. 4675–4685.

18. Rochette, L., et al., Functional roles of GDF15 in modulating microenvironment to promote carcinogenesis. Biochim Biophys Acta Mol Basis Dis, 2020. 1866(8): p. 165798.

19. Hoeres, T., et al., PD-1 signaling modulates interferon-γ production by Gamma Delta (γδ) T-Cells in response to leukemia. Oncoimmunology, 2019. 8(3): p. 1550618.

20. Garnier, L., et al., IFN-γ-dependent tumor-antigen cross-presentation by lymphatic endothelial cells promotes their killing by T cells and inhibits metastasis. Sci Adv, 2022. 8(23): p. eabl5162.

21. Prager, I. and C. Watzl, Mechanisms of natural killer cell-mediated cellular cytotoxicity. J Leukoc Biol, 2019. 105(6): p. 1319–1329.

22. Smyth, M.J., et al., Nature’s TRAIL--on a path to cancer immunotherapy. Immunity, 2003. 18(1): p. 1–6.

23. Ardiani, A., et al., Androgen deprivation therapy sensitizes prostate cancer cells to T-cell killing through androgen receptor dependent modulation of the apoptotic pathway. Oncotarget, 2014. 5(19): p. 9335–48.

24. Bonaterra, G.A., et al., Increased Density of Growth Differentiation Factor-15+ Immunoreactive M1/M2 Macrophages in Prostate Cancer of Different Gleason Scores Compared with Benign Prostate Hyperplasia. Cancers (Basel), 2022. 14(19).

25. Lee, N., et al., HLA-E is a major ligand for the natural killer inhibitory receptor CD94/NKG2A. Proc Natl Acad Sci U S A, 1998. 95(9): p. 5199–204.

26. Wang, X., H. Xiong, and Z. Ning, Implications of NKG2A in immunity and immune-mediated diseases. Front Immunol, 2022. 13: p. 960852.

27. Laccetti, A.L. and S.K. Subudhi, Immunotherapy for metastatic prostate cancer: immuno-cold or the tip of the iceberg? Curr Opin Urol, 2017. 27(6): p. 566–571.

28. Donahue, R.N., et al., Abstract 4901: Short-course enzalutamide reveals immune activating properties in patients with biochemically recurrent prostate cancer. Cancer Research, 2016. 76(14_Supplement): p. 4901–4901.

29. Portale, F. and D. Di Mitri, NK Cells in Cancer: Mechanisms of Dysfunction and Therapeutic Potential. Int J Mol Sci, 2023. 24(11).

30. Alspach, E., D.M. Lussier, and R.D. Schreiber, Interferon γ and Its Important Roles in Promoting and Inhibiting Spontaneous and Therapeutic Cancer Immunity. Cold Spring Harb Perspect Biol, 2019. 11(3).

31. Zingoni, A., et al., Cross-talk between activated human NK cells and CD4+ T cells via OX40-OX40 ligand interactions. J Immunol, 2004. 173(6): p. 3716–24.

32. Clavijo-Salomon, M.A., et al., Human NK cells prime inflammatory DC precursors to induce Tc17 differentiation. Blood Adv, 2020. 4(16): p. 3990–4006.

33. Herbst, R.S., et al., COAST: An Open-Label, Phase II, Multidrug Platform Study of Durvalumab Alone or in Combination With Oleclumab or Monalizumab in Patients With Unresectable, Stage III Non-Small-Cell Lung Cancer. J Clin Oncol, 2022. 40(29): p. 3383–3393.

34. Cascone, T., et al., Neoadjuvant Durvalumab Alone or Combined with Novel Immuno-Oncology Agents in Resectable Lung Cancer: The Phase II NeoCOAST Platform Trial. Cancer Discov, 2023. 13(11): p. 2394–2411.

35. Patel, S.P., et al., Phase 1/2 study of monalizumab plus durvalumab in patients with advanced solid tumors. J Immunother Cancer, 2024. 12(2).

36. Zorko, N., et al., Pan-cancer analysis of natural killer (NK) cell infiltration in human malignancies: Molecular features and clinical implications. Journal of Clinical Oncology, 2023. 41(16_suppl): p. 2563–2563.

37. Pasero, C., et al., Highly effective NK cells are associated with good prognosis in patients with metastatic prostate cancer. Oncotarget, 2015. 6(16): p. 14360–73.

38. Xu, L., et al., Inhibition of IL-6-JAK/Stat3 signaling in castration-resistant prostate cancer cells enhances the NK cell-mediated cytotoxicity via alteration of PD-L1/NKG2D ligand levels. Mol Oncol, 2018. 12(3): p. 269–286.

39. Wu, J., Could Harnessing Natural Killer Cell Activity Be a Promising Therapy for Prostate Cancer? Crit Rev Immunol, 2021. 41(2): p. 101–106.

40. André, P., et al., Anti-NKG2A mAb Is a Checkpoint Inhibitor that Promotes Anti-tumor Immunity by Unleashing Both T and NK Cells. Cell, 2018. 175(7): p. 1731–1743.e13.

41. Mahapatra, S., et al., High-resolution phenotyping identifies NK cell subsets that distinguish healthy children from adults. PLoS One, 2017. 12(8): p. e0181134.

42. André, P., et al., Differential regulation of killer cell Ig-like receptors and CD94 lectin-like dimers on NK and T lymphocytes from HIV-1-infected individuals. Eur J Immunol, 1999. 29(4): p. 1076–85.

43. de Kruijf, E.M., et al., HLA-E and HLA-G expression in classical HLA class I-negative tumors is of prognostic value for clinical outcome of early breast cancer patients. J Immunol, 2010. 185(12): p. 7452–9.

44. Seliger, B., et al., HLA-E expression and its clinical relevance in human renal cell carcinoma. Oncotarget, 2016. 7(41): p. 67360–67372.

45. Zeestraten, E.C., et al., Combined analysis of HLA class I, HLA-E and HLA-G predicts prognosis in colon cancer patients. Br J Cancer, 2014. 110(2): p. 459–68.

46. Borst, L., S.H. van der Burg, and T. van Hall, The NKG2A-HLA-E Axis as a Novel Checkpoint in the Tumor Microenvironment. Clin Cancer Res, 2020. 26(21): p. 5549–5556.

